# Functional Biomarkers *of Ex-vivo* Dental Caries Onset

**DOI:** 10.1101/2022.04.01.486588

**Authors:** Dina G. Moussa, Ashok K. Sharma, Tamer Mansour, Bruce Witthuhn, Jorge Perdigao, Joel D. Rudney, Conrado Aparicio, Andres Gomez

## Abstract

**Background:** The etiology of dental caries remains poorly understood. With the advent of next generation sequencing, a number of studies have focused on the microbial ecology of the disease. However, taxonomic associations with caries have not been consistent. Researchers have also pursued function-centric studies of the caries microbial communities aiming to identify consistently conserved functional pathways. A major question is whether changes in microbiome are a cause or a consequence of the disease. Thus, there is a critical need to define conserved functional biomarkers at the onset of dental caries.

**Methods:** Since it is unethical to induce carious lesions clinically, we developed an innovative longitudinal *ex-vivo* model integrated with the advanced non-invasive multiphoton second harmonic generation bioimaging to spot the very early signs of dental caries, combined with 16S rRNA short amplicon sequencing and liquid chromatography-mass spectrometry-based targeted metabolomics.

**Findings:** For the first time, we induced longitudinally-monitored caries lesions validated with the scanning electron microscope. Consequently, we spotted the caries onset and, associated to it, distinguished five differentiating metabolites - Lactate, Pyruvate, Dihydroxyacetone phosphate, Glyceraldehyde 3-phosphate (upregulated) and Fumarate (downregulated). Those metabolites co-occurred with certain bacterial taxa; *Streptococcus, Veillonella, Actinomyces, Porphyromonas, Fusobacterium*, and *Granulicatella*, regardless of the abundance of other taxa.

**Interpretation:** These findings are crucial for understanding the etiology and dynamics of dental caries, and devising targeted interventions to prevent disease progression.

**Funding:** The study was funded by the National Institute for Dental and Craniofacial Research of the National Institutes of Health and the University of Minnesota.

**Research in Context:** *Evidence before this study:* Studies have shown that dental caries, tooth decay, occurs as a result of disruptive imbalance in the oral ecosystem. Excessive dietary intake of fermentable carbohydrates is a critical contributor to disease progression by promoting bacterial production of acids, which shifts the microbial community to an imbalanced and a less diverse one. Studies have also shown that microbial associations with caries have not been consistent while their functions are relatively conserved across individuals. Still, the specific microbial functions associated with the dental caries onset is still unknown due to its infeasible clinical diagnosis.

*Added value of this study:* This study applied a novel longitudinal *ex-vivo* model, integrated with advanced non-invasive bioimaging, for experimental dental caries induction. This model enabled the detection of the exact onset of the disease, which is undetected clinically. Then, the microbial communities accompanying the caries onset were analyzed for their microbial composition and metabolic functions in comparison to normal conditions. Our study identified five metabolites differentiating the caries onset. Further, we investigated the co-occurrence of these metabolic biomarkers with certain oral bacteria.

*Implications of all the available evidence:* Our study provides carefully validated evidence for biomarkers of the dental caries onset. These data are critical for early diagnostics and development of timely intervention strategies to interfere with the disease progression that otherwise requires invasive and costly health care expenses. Moreover, our data open new avenues for developing therapeutics to neutralize the identified metabolic biomarkers or target the accountable bacteria for caries prevention.

## Introduction

Although dental caries remains the most common chronic disease globally, affecting more than 95% of adults (1, 2) (3-5) there is a significant gap in our understanding of its exact underlying pathogenesis. Dental caries can be considered as the outcome of dysbiotic changes in the biofilm community of supragingival dental plaque (6, 7). Demineralized carious lesions occur as the cumulative outcome of repeated shifts towards a less diverse microbiota that produces and tolerates a low pH (8) (9, 10). This disruptive imbalance in the oral ecosystem is often caused by excessive dietary intake of readily fermentable carbohydrates (11, 12). In metagenomic and marker gene studies, caries-associated communities are typically less diverse than healthy supragingival plaque. However, those dysbiotic communities still display considerable taxonomic diversity between affected individuals and, notably, caries associations have not been consistent between studies (13-17). On the other hand, a consensus has been reached that dental caries is a community-scale metabolic disorder (7, 18, 19),(20) (21) and despite the large variation in the microbial community structure, conserved metabolic pathways exist across individuals for supragingival plaque (22). Thus, there is a critical need to define functional biomarkers of dysbiosis that are less dependent on taxonomy (21, 23, 24) to better understand the disease etiology and pathogenesis. ***Correspondingly, our first hypothesis is*** that supragingival plaque microbial communities undergo conserved changes in metabolism during caries-inducing conditions, regardless of taxonomic assortment.

Another layer of complication is that caries is a dynamic and progressive disease associated with continuous changes of the community compositional profile (21). It remains unclear how these progressive changes impact microbiota function early in the disease course. Accordingly, the analysis of the functional changes associated with the transition from health to carious lesions is critical to determine etiological factors of this disease. Unlike the reversible early periodontal diseases, such as gingivitis, that can be induced by refraining the patients from brushing over a three-week experimental period (25), inducing clinical early carious lesions is not possible, which hinders the analysis of the associated microbial functions in the transition from health to disease. Alternatively, examining a massively large numbers of healthy and disease-associated microbial communities could allow the characterization of their corresponding common features. However, this would represent an unfeasible experimental approach in addition to its limitation to spot the lesions in preclinical stages or at the early onset. Thus, *ex-vivo* models to induce carious lesions emerge as the most clinically-relevant alternative approach. These models have been optimized to obtain reproducible biofilms with taxonomic and metabolic diversity approaching that of the human oral microbiome (26, 27). The successful implementation of these *ex-vivo* models requires a reliable assessment of the different stages during progression of the disease. However, access to non-invasive monitoring systems that are sensitive enough to detect the early dental changes at the disease onset and can be implemented to longitudinally monitoring the progression of lesions has been a technological hurdle.

With the advent of multiphoton second harmonic generation (MP-SHG) imaging technology, early recognition of *ex-vivo* induced dental caries has been enabled through non-invasive and label-free monitoring of the subtle changes in both mineralized and collagenous phases comprising the dental hard tissues (28, 29). MP-SHG is based on a pulsed near-infrared laser allowing the excitation of the biological samples to penetration depths inaccessible to 1-photon conventional confocal microscopy (30). The MP component, that collects the signals for the mineral phase (enamel), strongly suppresses the background signal, optimizes the signal-to-noise ratio, and reduces the phototoxicity to the focal region for high resolution optical sectioning (28, 31). Meanwhile, the SHG that depends on the polarization, orientation, and symmetry properties of collagen chiral molecules offers novel opportunities to investigate the three-dimensional structure of the label-free collagen molecules within biological tissues (29, 32, 33). Slimani et al 2018 showed that SHG imaging of dentin produced extremely accurate signals for detecting early stages of dental caries as dentin is particularly efficient in producing SHG (29) because of the presence of self-assembled fibers of collagen type I; >90% of the organic matrix (34).

***Herein, we also hypothesize that*** the longitudinal analysis of the genomic diversity and metabolic profiles of supragingival microbiota associated with *ex-vivo* dental caries onset and after progressing to overt lesions enables the detection of caries etiological factors. To our knowledge, this will be the first study integrating advanced bioimaging, microbial ecology and metabolomics analyses to study the etiology of dental caries implementing an *ex-vivo* caries-induction model with reproducible and taxonomically diverse oral microcosm biofilms. Our ultimate goal is to capture the conserved functional biomarkers accompanying the induced caries onset to determine the etiology of the disease at the early, otherwise clinically undiagnosable stage. Better understanding of the microbial functions associated with early onset of carious lesions is critical to develop targeted prophylactic approaches to prevent the progression of this widespread disease and to reduce the associated massive health care expenditures.

## Material and Methods

### Microcosm supragingival plaque biofilm

In this study, we used the reproducible taxonomically diverse oral microcosm biofilm model of dental caries developed by Rudney et al., 2012 (27). Briefly, 11 supragingival dental plaque samples from high caries risk subjects were collected at the University of Minnesota (Institutional Review Board #1403M48865). All subjects were in good general health and had not taken antibiotics within 3 months of plaque sampling. The microcosm biofilms were grown on hydroxyapatite (HA) discs in CDC biofilm reactor without (NS) or with sucrose (WS) to simulate the oral conditions in the flow of saliva and dietary carbohydrates across biofilm surfaces as described in (27). Each pair of NS and WS microcosm was grown from a single plaque inoculum mimicking health and sucrose-induced dysbiotic models that evolved from the same patient.

### Teeth preparation

A total of 30 sound extracted human third molars were selected from a pool of unidentified extracted teeth previously obtained as surgical waste from oral surgery clinics, that is exempt from IRB review, at the University of Minnesota and in the Minneapolis/Saint Paul metropolitan area (35). They were screened visually, tactilely by explorers, and by magnifying stereomicroscope (MVX10, Olympus, Tokyo, Japan) for any early carious lesions. 14 teeth were selected to be used as substrates to grow biofilms after slicing. The roots as well as the proximal enamel sides were sectioned using a diamond saw (Isomet™, Buehler, Lake Bluff, IL, USA) and discarded. Coronal proximal slices were then sectioned occluso-cervically from the mesial and distal sides of each tooth. Each resultant slice included enamel and dentin tissues. The sliced specimens were then ground to 0.2 mm, polished with 320-, 600-, and 1200-grit Si-C papers using a polishing machine (Ecomet 3, Buehler, IL, USA), and ultrasonicated in a water bath for 20 min (36).

### Pre-inoculation screening of dental specimens with stereomicroscopy and MP-SHG

Fourteen pairs of polished teeth specimens were screened initially with the stereomicroscope then analyzed at high magnification with non-invasive MP-SHG advanced microscopy to distinguish any early lesions or structural anomalies according to the protocol we previously developed for dental hard tissue bioimaging (36, 37). MP-SHG collects the MP fluorescence signals emitted from the mineralized phase, mainly in enamel, and the SHG signals emitted from the collagen network, exclusively in dentin, non-linearly and *via* 2 separate channels without labelling and in fully-hydrated conditions (32). MP-SHG images were acquired using a Leica SP5 confocal laser scanning microscope (Leica Microsystems CMS GmbH, Am Friedensplatz 3, D-68165 Mannheim, Germany). Excitation of the samples was performed with a Spectra-Physics 15W Mai Tai DeepSee tunable IR laser tuned to 870 nm. Non-descanned highly sensitive GaAsP detectors were used to allow visualization of multiple fluorophores located deep within the specimen. The IR laser was operated in pulsed mode and was focused onto the sample using a 25x NA0.95 water-immersion objective with a 2.4 mm working distance suitable for deep imaging. The SHG signal was collected at 400-450 nm using a spectral HyD PMT and the multiphoton fluorescence was collected at 450-700 nm using a spectral PMT with a maximally open pinhole set to 10.74 airy units. The samples were scanned in 512×512 pixel frames (0.714×0.714 µm pixel size) and 50 µm of dentin thickness was sampled with a 0.49 µm step size in Z. Images were compiled and analyzed with FIJI software (Fiji Is Just) ImageJ, version: 2.1.0/1.53c https://imagej.net/Contributors (38).

### Caries onset induction

11 pairs of microcosm biofilms, NS and WS culturing conditions, were grown on the 11 pairs of the proximal coronal sections of human teeth. Each pair of either microcosm biofilms or teeth specimens was originated from a single patient (Figure 1). The teeth specimens were disinfected by immersion in 70% ethanol for 5 min and left to dry for 10 min in a biosafety cabinet then loaded in 24 suspension culture well plate to be inoculated. Overnight cultures of NS and WS microcosms were grown in basal mucin medium (BMM) (39) without or with 5% sucrose supplementation, respectively. BMM is a complex medium that was developed as saliva analog and has been used successfully in previous oral microcosm models containing hog gastric mucin as the primary source of carbohydrate (Supplemental Table 1) (23, 27). The inoculum was adjusted at 1×10^6^ cell/ml and the inoculated samples were incubated aerobically at body temperature under continuous shaking to simulate the ecological succession of supragingival plaque. The WS samples were subjected to sucrose bath pattern of 5h/day, hence the samples were switched between the BMM media with and without sucrose for periods of 5 and 19 h, respectively where the inoculation was consistently with the same WS microcosm. The pH was monitored at intervals up to 48 h where the media was replenished (Supplemental Table 2). New inoculation was introduced every 96 h. Based on results from our preceding pilot and corresponding to the development of incipient lesions, 88 h was set as the 1^st^ time point (1^st^ phase – caries onset) at which the experiment was paused for detaching biofilms.

**Figure 1.**
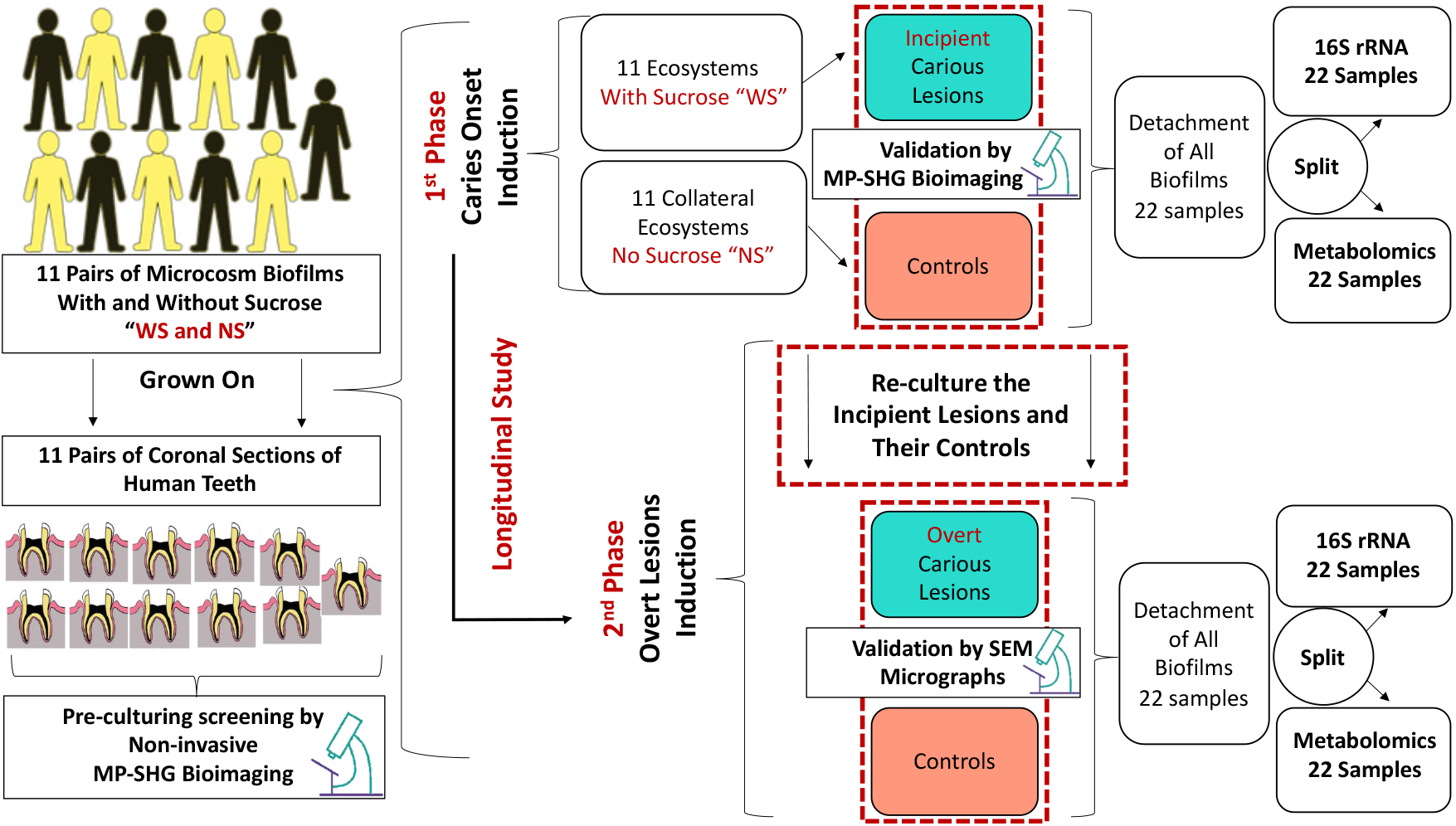
Schematic diagram depicting the study design and workflow for identifying bacterial and metabolites changes associated with dental caries onset and progressive overt lesions. Eleven pairs of supragingival plaque microcosms were developed in normal and dysbiotic conditions, without (NS) and with sucrose (WS) and grown on 11 pairs of human teeth slices. Each pair of microcosm was originated from a single individual and each pair of teeth specimens was sliced from a single tooth. All the specimens were screened at high magnification with the label-free and non-invasive Multiphoton-Second Harmonic Generation microscopy (MP-SHG) to discard the specimens with any potential lesions before starting the inoculation. The dental caries was induced *ex-vivo* and the associated biofilms were analyzed at two longitudinal phases, the first and second phases correspond to the caries onset and overt lesions, respectively. The caries onset was validated with MP-SHG before proceeding with inducing progressive overt lesions. Induced caries onset as well as overt lesions were further validated individually with Scanning Electron Microscopy (SEM). The biofilms associated with all samples at the 2 phases were detached and split-divided for16S rRNA genomic analysis and targeted central carbon metabolomic analysis. Along the manuscript, the red and green shades denote the samples grown in control and dysbiotic conditions, respectively.

### Detaching biofilms

The media was gently aspirated and the specimens were washed with sterile 10 mM ammonium bicarbonate. Each sample was then immersed in 1.5 ml of cold 10 mM ammonium bicarbonate (Sigma-Aldrich, Burlington, MA), bath sonicated, to minimize heating, for 30 min followed by brief vortexing for detaching biofilms (40). Ammonium bicarbonate was used due to its compatibility with the liquid chromatography-mass spectrometry and moderate pH buffering (41). The detached biofilms were transferred to 1.5 ml low retention microcentrifuge tubes and split-divided for extracting DNA and metabolites separately for the 1^st^ time point analysis.

### Stereomicroscopy and MP-SHG monitoring of the caries onset

All samples were re-screened in their well-plates for any visualized early lesions using the stereomicroscope. Then, samples were loaded fully hydrated in sterile and adhesive silicon isolators (Press-to-Seal™, Thermo Fischer Scientific) for cellular imaging to avoid any cross contamination during MP-SHG monitoring of early lesions. All imaging parameters were set as mentioned above in the pre-inoculation screening and throughout the study.

### Overt caries lesions induction

Biofilms were regrown on same samples using same microcosms under same conditions as mentioned above till overt lesion were visualized by naked eyes after 17 days. All biofilms were detached as detailed above and split-divided for extracting DNA and metabolites for the 2^nd^ time point analysis (2^nd^ phase – overt lesions). The overt lesions were visualized with the stereomicroscope and examined with MP-SHG for ultrastructural characterization.

### Scanning electron microscopy characterization of the induced lesions

Scanning electron microscope (SEM) analysis was conducted to validate if *ex-vivo* induced lesions show structural changes similar to the clinical carious lesions at the dentin-enamel junction (DEJ) area for the induced incipient or/and overt lesions. Slices of sound teeth were used as a control after a mild etching, pH=1.95, with 25 % polyacrylic acid (50,000 wt.%) in H_2_O to remove the smear layer with minimal associated demineralization. These specifications were reached after a few pilots testing a range of different polyacrylic acid concentrations (10%-25%), different molecular weights (2000-50,000 wt.%), different application time (5, 7, 10, and 15 s), and different pH (1.5-3.2).

The samples were prepared following the optimized protocols developed by Perdigão et al as detailed in (43). Samples were fixed in 2.5% glutaraldehyde with 2% paraformaldehyde in 0.1 M cacodylate buffer (Electron Microscope Sciences, Hatfield, PA) at pH=7.4 for 12 h at 4°C, rinsed in 0.2 M cacodylate buffer at pH=7.4 for three consecutive periods of 20 min each, and rinsed with distilled water for 1 min. Then, the samples were dehydrated in ascending grades of ethanol: 25%, 50%, 75% ethanol for 15 min each, 95% ethanol for 30 min, and 100% ethanol for 60 min. Under the hood, the samples were immersed in 50/50 100% ethanol/ hexamethyldisilazane (HDMS) for 2 min. Then, the samples were transferred to absolute HDMS for 10 min and were let to dry in a desiccator with silica for 24 h. The HDMS desiccation method was used to replace the critical point drying step aiming to better preserve the collagen network and the micro-porosity of the demineralized dentin surface (43). The samples were mounted on aluminum stubs with a double-sided carbon tape and the periphery was painted with silver ink to avoid over-flowing and disturbing the imaging area of interest as well as enhancing the electron conductivity. The samples were then coated with 5 nm iridium (EM ACE600, Leica Microsystems Inc., Buffalo Groove, IL) and observed with a Hitachi S-4700 field-emission scanning electron microscope (Hitachi, Tokyo, Japan) at an accelerating voltage of 5-10 kV and a working distance of 12-14 mm.

### DNA extraction

As mentioned above, the detached biofilms from all samples at the two time points, 44 samples in total other than the controls (control media and control teeth slices), were split-divided for microbiome and metabolome analyses. Genomic DNA was extracted following the protocol amended from Epicentre MasterPure™ DNA Purification Kit (http://homings.forsyth.org/DNA%20Isolation%20Protocol.pdf). The DNA yield was assessed with a NanoDrop™ 2000 spectrophotometer. The yield extracted from WS biofilms, i.e., grown in cariogenic conditions, was consistently higher compared to NS biofilms (Supplemental Table 3). Samples were submitted to the University of Minnesota Genomics Center for dual-indexed amplicon sequencing of 16S rRNA bacterial gene (V4 region) on the Illumina MiSeq platform (2x 300PE) (primers 515F (5’-GTGCCAGCMGCCGCGGTAA-3’) and 806R (5’-GGACTACHVGGGTWTCTAAT-3’)).

### Metabolites extraction

Efficient extraction of the metabolite class of interest is essential to the success of targeted metabolomics approaches. This is because, unlike untargeted metabolomics, we focus only on a subset of metabolites for downstream analyses. The extraction method was tailored to the physico-chemical properties of the byproducts of central carbon metabolism, excluding other components such as proteins (44). Briefly, the half of detached biofilms assigned for metabolites extraction was pelleted at 5000x g for 10 min at 4 °C, then resuspended in a mix of 50 µl of 10 mM ammonium bicarbonate and 50 µl of methanol. Three freeze-thaw cycles were conducted where samples were frozen at −80 °C for 15 min and then, thawed in a water bath at room temperature for 10 min with 1 min vigorous vortexing in-between cycles. Then, 4 volumes of chilled 90/10 methanol/acetone were added to each sample and vortexed at high speed for 1 min to denature the proteins. Samples were then incubated at −10°C for 15 min and centrifuged at 13,000 x g for 15 min at 4°C. Supernatants were carefully transferred to new microfuge tubes avoiding the soft pellets. The samples were dried by evaporation under a stream of inert nitrogen gas and reconstituted in 75 µl of 5% acetonitrile, 0.1% formic acid. To resuspend any particulates, the samples were vortexed and then centrifuged at 13,000 x g for 5 min at 4 **°**C. Samples were stored at −80**°**C until used for mass spectrometry.

### 16S rRNA sequencing

Raw reads were processed to remove primers and low-quality base pairs (q<30) using cutadapt (45) and fastx_toolkit (46). Processed high quality sequences were further processed using the DADA2 plugin (47) within Qiime2 (46) to generate Amplicon Sequence Variants (ASVs). For taxonomic assignment of these ASVs, reference sequences (clustered at 99% sequence identity) from the Greengenes database, v13_8 (48) were downloaded and used for training the naïve Bayes classifier, using the feature-classifier fit-classifier-naive-bayes function of Qiime2. This trained classifier was then used for assigning taxonomy for each ASV detected using the feature-classifier classify-sklearn function of Qiime2. Taxa abundances at the ASV level were then used for downstream statistical analysis.

### Targeted analysis of the central carbon metabolism (Liquid Chromatography-Mass Spectrometry “LC-MS”)

Central carbon metabolites were analyzed using the Selective Reaction Monitoring (SRM). Samples (10 µl) for SRM analysis were subjected to separation using a Shimadzu system, coupled to an analytical SeQuant ZIC-pHILIC (150 mm x 4.6 mm at 30 °C connected to the Applied Biosystem 5500 iontrap) and fitted with a turbo V electrospray source run in negative mode with declustering potential and collision energies as listed in table (Supplemental Table 4). The samples were subjected to a linear gradient of A: 75% Acetonitrile, B: 20% acetonitrile, 10 mM ammonium acetate for 22 min at a column flow rate of 400 µl/min. The column was cleared for 2 min with 60% B and then equilibrated to buffer A for 10 min.

Transitions were monitored as shown in (Supplemental Table 4) and were then established using the instrument’s compound optimization mode with direct injection for each compound. The data were analyzed using MultiQuant™ (ABI Sciex Framingham, MA), which provided the peak area. A standard curve was constructed from picomole to nanomole in 10 µl. Samples were run in duplicate and concentrations determined from the standard curve.

### Bioinformatics and statistical analysis

#### 16S rRNA analyses

Statistical analyses of microbiome data were performed using the R statistical interface, version 4.0.2 (49). ASV tables were filtered using the R labdsv package (50) to remove ASVs that were likely to be sequencing artifacts due to their presence at extremely low frequencies or only in 3 or fewer samples. Alpha diversity analyses, beta diversity using Bray-Curtis distances and permutational multivariate analysis of variance (PERMANOVA) were performed using the vegan package (51). Principal coordinate analyses (PCoA) based on Bray-Curtis or Euclidean distances, were created using the ape package (52). Discriminating taxonomic features were identified based on fold changes and *p*-values calculated using the DESeq package (53), and by indicator species analyses using the labdsv package (54).

#### Metabolome analyses

Metabolomic analyses were performed using **MetaboAnalyst 5.0** https://www.metaboanalyst.ca/ (55). Briefly, LC-MS data were normalized using Log transformation and Pareto scaling. Partial least squares discriminant analyses (PLS-DA) and variable importance in projection (VIP) scores were used to explore the extent to which the metabolite data predicted phenotypes of interest and to identify metabolites that distinguished each phenotype. PLS-DA analyses were validated through permutation tests, including Q2 and R2 statistics. Metabolomic data were also visualized *via* clustering analyses (Euclidean Distance and weighted Averages plotted on heatmaps) and using PCoA, in tandem with PERMANOVA as described with the microbiome data. Associations between metabolomic and microbiome data were performed using Procrustes and Mantel tests in the vegan package of R (56) and mmvec (57).

The false discovery rate (FDR) was applied for multiple comparisons correction of all microbial and metabolite biomarkers to limit false positives while maximizing power. The FDR creates a balance between the number of true and false positives that is automatically calibrated (*q* value) with each tested feature (58). The significance cut-off of the FDR adjusted *p*-value (or *q* value) was set at 0.05. The sample size was outlined based on the developed 11 pairs of the reproducible oral microcosm biofilm model for dental caries by Rudney et al., 2012 (27).

## Results

In this study, we analyzed the 16S rRNA bacterial gene, targeted central carbon metabolites *via* LC-MS, and their interrelationships on a supragingival dental plaque microcosm grown *ex-vivo* in normal and cariogenic conditions (Figure 1).

### Pre-inoculation screening of dental specimens with stereomicroscopy and MP-SHG

MP-SHG and stereomicroscopy representative images of pre-inoculated teeth slices are shown in (Figure 2 and 3) and a set of images for all screened validated samples are depicted in (Supplemental Figure 1). Two of the screened specimens showed attenuated signals, which suggested the presence of potential subclinical lesions (28, 29). Accordingly, these two specimens and their parent teeth were excluded from the study.

**Figure 2.**
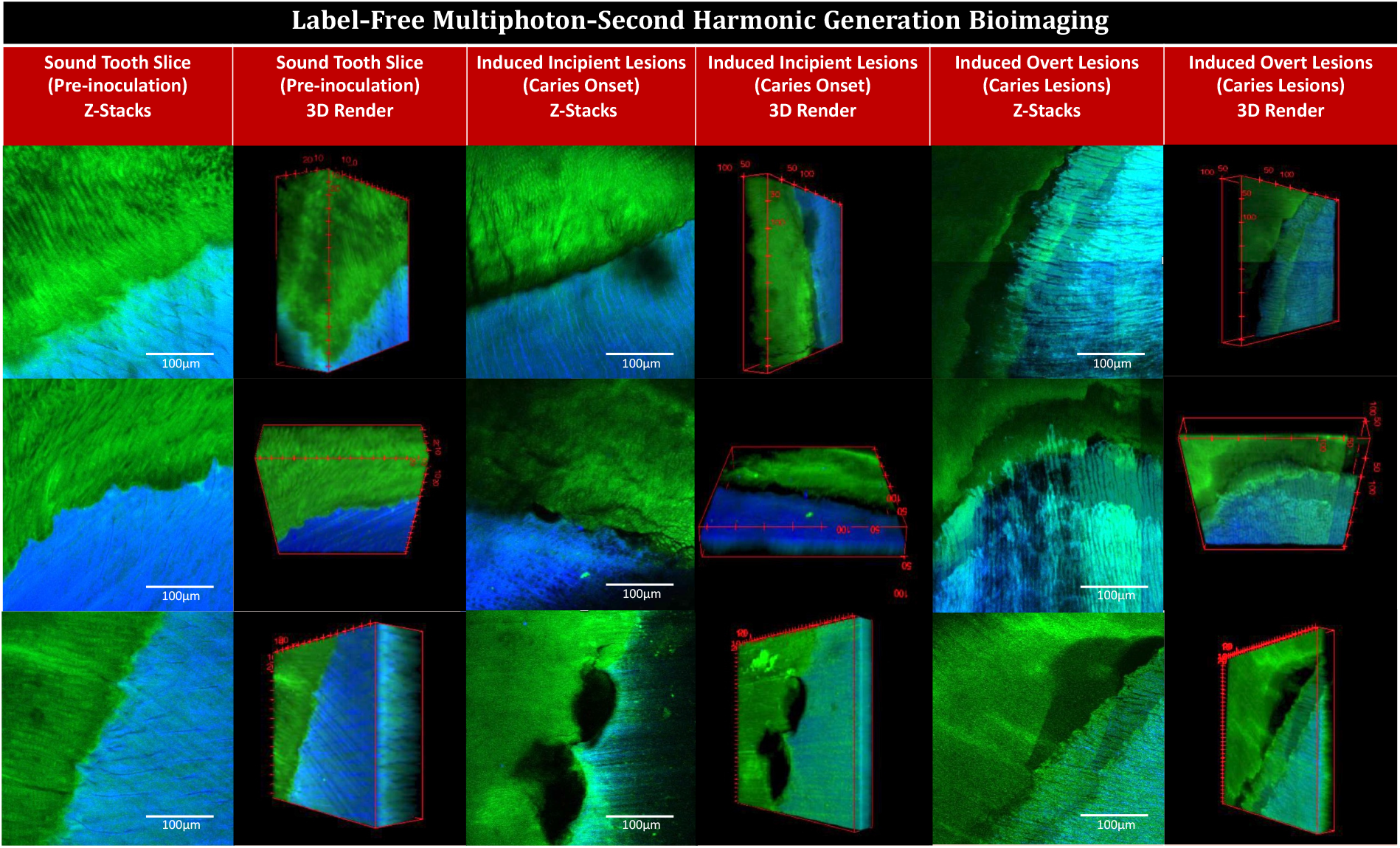
Multiphoton-second harmonic generation (MP-SHG) bio-imaging examination of the dentin-enamel junction (DEJ) area before and along the course of *ex-vivo* dental caries induction. The green signals show the MP autofluorescence emitted from the mineralized phase (mainly enamel). The blue signals show the SHG exclusively emitted from the collagen network of dentin. The columns show the maximum intensity projection of the acquired Z-stacks and the 3D renders of sound teeth slices (pre-inoculation), induced incipient caries lesions (caries onset), and induced overt caries lesions; the videos are provided in Supplemental Videos 1. The rows show three representative samples; all tested samples are displayed in Supplement Figures 1 and 3.

**Figure 3.**
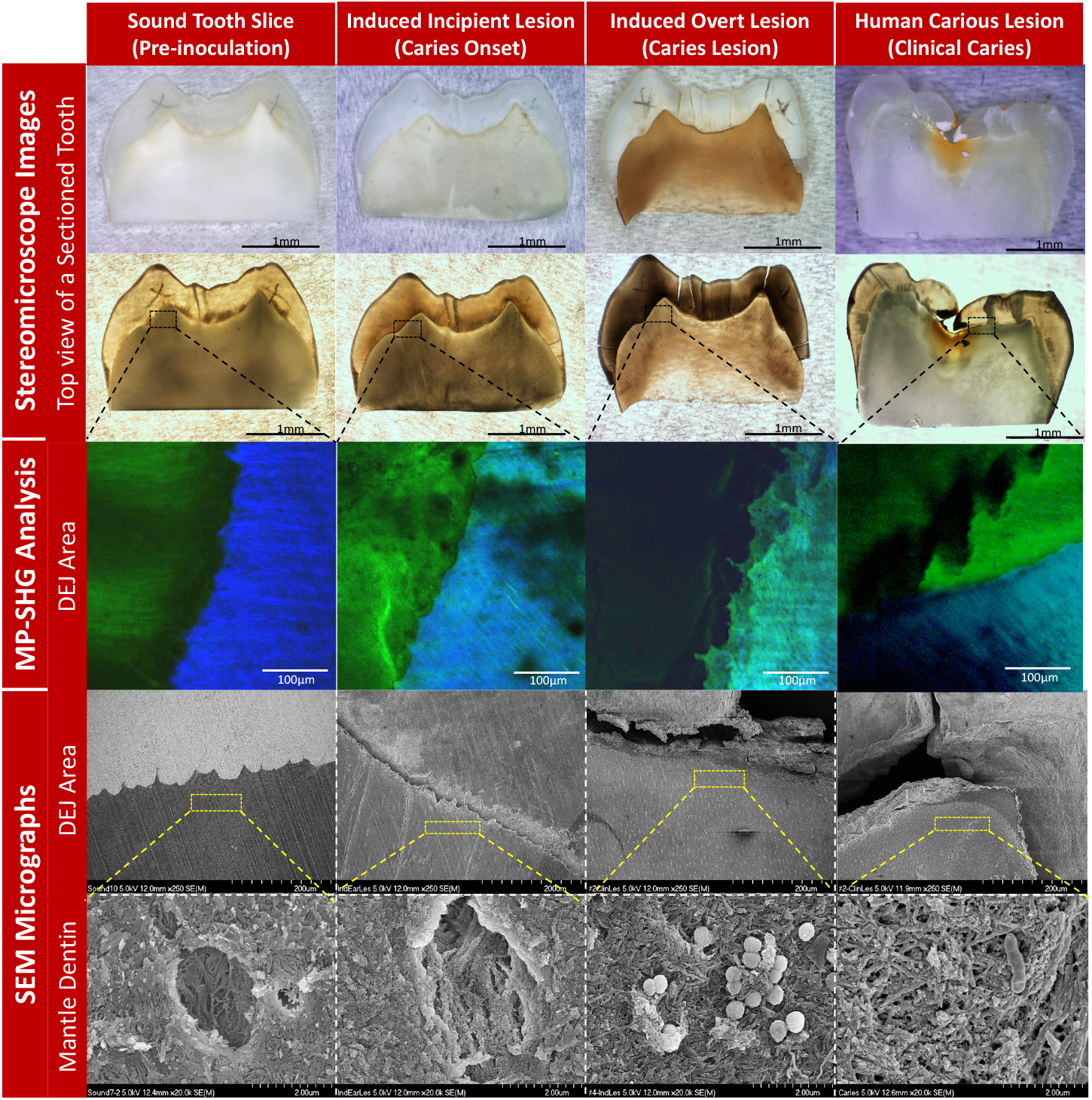
Characterization of the stages of induced *ex-vivo* caries lesion in comparison to the clinical caries lesion. The columns from left to right show the pre-inoculation stage, induced caries onset, induced overt caries lesion, and clinical caries lesion. The rows from top to bottom show the characterization methods with stereomicroscopy, Multiphoton-second harmonic generation microscopy (MP-SHG) and Scanning Electron Microscopy (SEM). The stereomicroscope images show the overall changes of the teeth specimens in reflection and transmission light modes. The specific dentin-enamel junction (DEJ) areas assigned for ultrastructural characterization are outlined with black dashed boxes. The MP-SHG examination shows the degradation of the enamel and dentin around the DEJ and spotted the early lesion in the mantle dentin zone, just beneath the DEJ. The SEM further characterized the ultrastructural changes of the mantle dentin associated with the *ex-vivo* caries induction compared to a clinical caries lesion. The micrographs showed the gradual degradation of the peritubular dentin and disorganization of the collagen fibers along the cariogenesis course; portraying the structural similarities between the induced overt lesions and clinical caries developed in the same dentin area.

### Induction of ex-vivo caries lesions

Caries lesions were induced by inoculating 11 pairs of human teeth slices with eleven pairs of supragingival plaque microcosms, developed in normal and dysbiotic conditions, without (NS) and with sucrose (WS) supplementation, respectively. In the first 3 h after inoculation, the pH of all bacterial cultures, NS and WS, were ∼ 6.5-7. Afterwards, the acidity of the WS cultures increased significantly and the pH dropped between 3-4 while NS cultures remain at the range of pH=7-8 (Supplemental Table 2). The same pattern of pH changes was observed with the periodic media change (every 48h) and with the periodic inoculation (every 96h).

### MP-SHG characterization of the induced caries lesions

After 88h of incubation, initial spots of disintegration were observed using MP-SHG microscopy across WS samples, particularly around the dentin-enamel junction (DEJ) (Figure 2, Supplemental Figure 1, and Supplemental Videos 1). None of these incipient lesions were recognized with either the naked eye or stereomicroscopy (Figure 3 and Supplemental Figure 2). After 17 days of incubation, overt brownish soft lesions were detected in dentin below the DEJ area in the WS samples (Figure 3 and Supplemental Figure 2). The MP-SHG characterization revealed a distinct band of disintegrated dentin just below the DEJ area (Figure 2, Supplemental Figure 3, and Supplemental Videos 2). The collateral specimens that were sliced from same teeth and were inoculated with the same microcosm but with no sucrose supplementation (NS), did not show similar areas of disintegration or discoloration at any stage of analysis (Supplemental Figure 2).

### SEM characterization of the induced caries lesions

At a low magnification (250 X), SEM micrographs showed signs of demineralization and degradation at the DEJ of the induced incipient lesions compared to the intact sound controls (Figure 3). Complete separation between enamel and dentin at the DEJ was observed in the induced overt lesion with substantial signs of demineralization, which appeared notably similar to that of actual clinical lesions (Figure 3). Observation at high magnification of the mantle dentin, 30 µm below the DEJ, revealed that the dentinal tubules were surrounded by a thickened collar of peritubular dentin in the sound controls and the collagen fibers had sound organized arrangements (Figure 3). However, in the induced incipient lesions, the peritubular dentin collar was partially demineralized with areas of discontinuation along the perimeter of the tubules, accompanied with collagen fibers disorganization (Figure 3). Induced overt lesions and clinical lesions displayed complete demineralization and disorganization of the peritubular dentin. Collagen fibers had also signs of denaturation as the collagen banding pattern was lost (Figure 3). Similar demineralization patterns were observed for incipient and overt induced lesions at far distances (83µm and 272 µm) from the DEJ but less progressive (Supplemental Figure 4).

### 16S rRNA sequencing

We assessed beta and alpha bacterial diversity within the 16S rRNA data sets of supragingival plaque microcosms grown in dysbiotic (cariogenic, WS) and non-dysbiotic (control, NS) conditions at the caries onset (TimePoint1 “T1”), and after progression to overt caries lesions (TimePoint2 “T2”) (Figure 4). Beta diversity analyses (Bray-Curtis dissimilarity) showed distinct clustering, mainly according to time points sampled (PCoA axis 1, 45% of variation), followed by condition (PCoA axis 2, 16%) (Figure 4a). That is, overall shifts in bacterial community composition were more influenced by temporal scale rather than by treatment, where the most distinct and tightly clustered groups were WS_T2 followed by NS_T2; these groups represent the tested second time points of test and control groups (Figure 4a). Nonetheless, alpha diversity (Number of ASVs, Shannon and Simpson indices) did not show significant differences among the tested groups (Figure 4b). PERMANOVA confirms significant differences in microbiome composition (F-model: 10.83, 16.87, 2,71; R^2^: 0.15, 0.24, 0.038; and *P*-value: 0.001, 0.001, 0.012) for treatment, time points and their interactions respectively (Supplemental Figure 5c).

**Figure 4.**
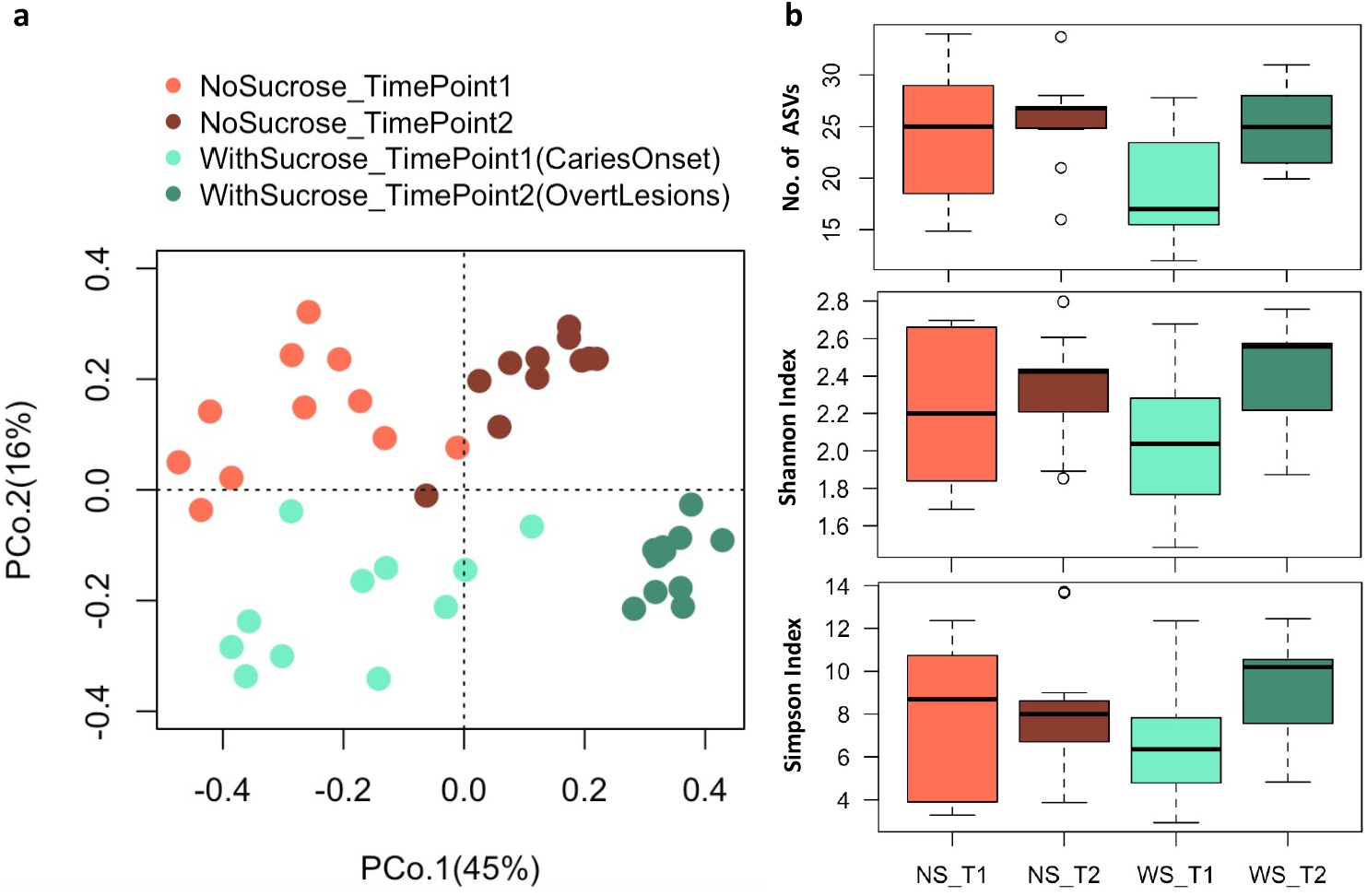
Beta and alpha diversity of supragingival plaque microcosms grown in dysbiotic (cariogenic) and non-dysbiotic (control) conditions at the *ex-vivo* dental caries onset and after progression. **A)** Principal coordinates analysis (PCoA) of 16S rRNA sequencing reads of all tested samples. The red and green shades denote the samples grown in control and dysbiotic conditions respectively, where the lighter tones refer to the first time point and the darker tones refer to the second time point of analysis. 16S rRNA sequencing reads were categorized into distinct amplicon sequence variants (ASVs) using standard QIIME scripts. Beta-diversity between samples was measured through a Bray-Curtis dissimilarity analysis based on relative abundance of ASVs. The percent of variability accounted for by each axis is indicated. **B)** Differences in alpha diversity between dysbiotic and non-dysbiotic samples at each time-point were measured as number of amplicon sequence variants “No. of ASVs”, Shannon, and Simpson indices. No statistical differences were found among the alpha diversity indices of the tested groups. (T1, T2) stand for (Time Point 1-1^st^ phase/caries onset, Time Point 2-2^nd^ phase/overt lesions) and (WS, NS) stand for (With Sucrose, No Sucrose).

The relative abundance of the bacterial taxa (ASVs) characterizing each of the tested groups was visualized on volcano plots, showing fold changes and significance according to time point sampled and treatments using pairwise comparisons (Figure 5a and Supplemental Table 5 and 6). Few taxa showed significant changes timewise. For the NS groups, *Enterobacteriaceae* and *Atopobium* showed significant increase at T2 while one of *Streptococcus* ASV showed significant increase at T1. Likewise, the WS_T2 group showed a significant increase in *Corynebacterium* compared to NS_T2. Only a ASV of the genus *Bacillus* was detected significantly abundant in WS_T1 (Figure 5a and Supplemental Table 5). Sucrose-supplementation shifted several taxa consistently and significantly at any given time point, compared to the number of taxa observed with no sucrose groups, after the *p*-value was adjusted for the FDR.

**Figure 5.**
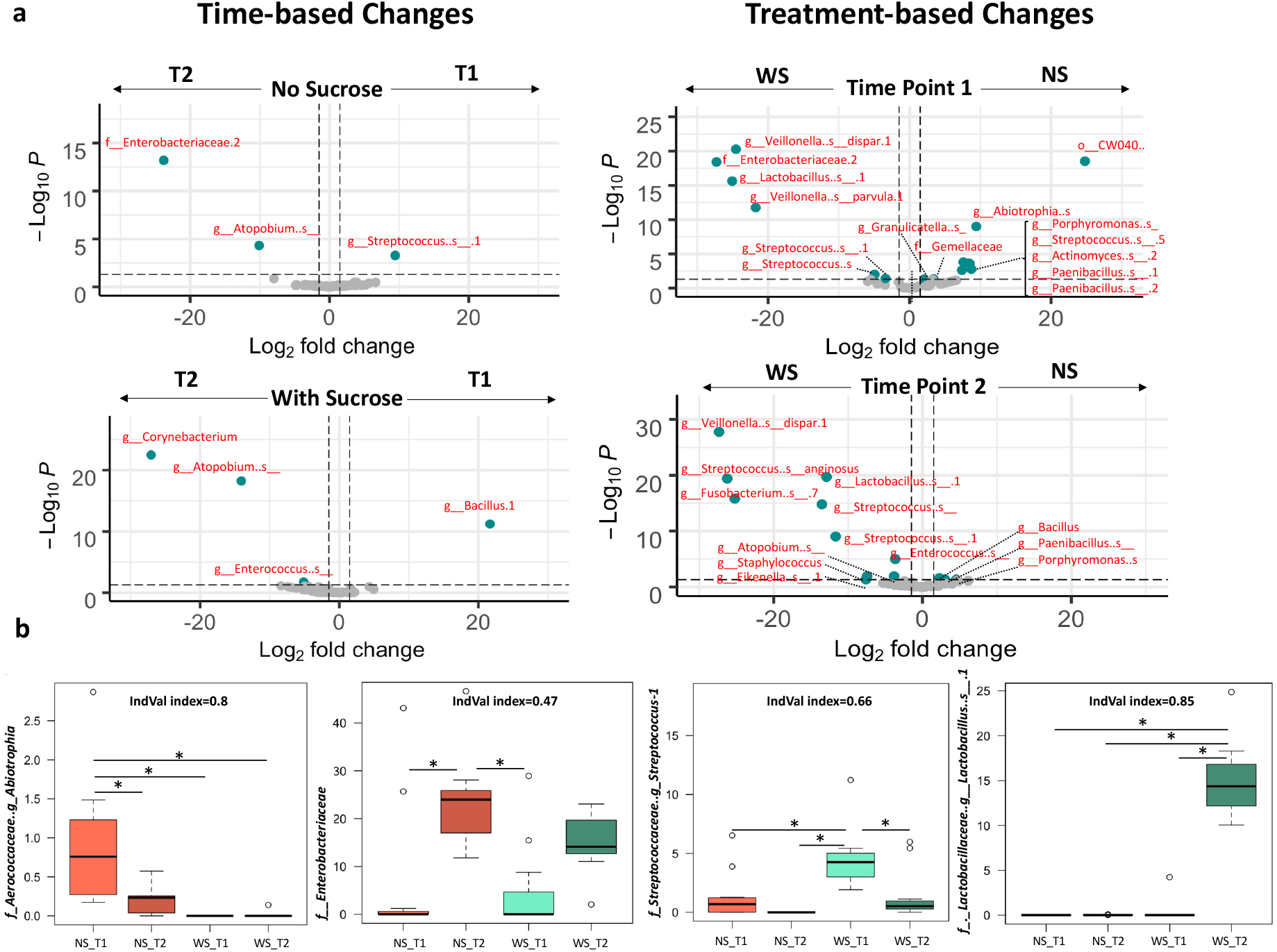
Differential relative abundance analysis of the supragingival plaque bacterial taxa associated with dysbiotic (cariogenic) and non-dysbiotic (control) conditions at the *ex-vivo* dental caries onset and after progression to over lesions. **A)** Volcano plots display the fold changes in relative abundance (log 2) on the x axes between the first and second time points (caries onset and overt lesion) on left (time-based changes) and between dysbiotic and non-dysbiotic conditions on right (treatment-based changes). The dashed vertical lines denote a 2-fold change in relative abundance (log_2_2 = 1). The *y* axes display the −log10 of the adjusted *p* values (*q values*) of the test statistic. The dashed horizontal line corresponds to the *q value* of 0.05. The green points that appear outside the enclosed dashed box formed by the *x* and *y* axis intercepts are points that show both significant and proportionally large shifts in relative abundance. The gray points below the significance threshold denote non-significant shifts in relative abundance. A full list of taxa showing significant shifts in relative abundance is provided in Figure5-table supplement 1 and 2. **B)** Indicator value “IndVal” analysis-based box plots showing the most abundant taxa associated with each group. The red and green shades denote the samples grown in control and dysbiotic conditions respectively, where the lighter tones refer to the first time point and the darker tones refer to the second time point of analysis. The detailed analysis of IndVal index showing the full list of taxa associated with each group is provided in Figure5-table supplement 3. (T1, T2) stands for (Time Point 1-1^st^ phase/caries onset, Time Point 2-2^nd^ phase/overt lesions) and (WS, NS) stands for (With Sucrose, No Sucrose).

WS_T1 displayed a significant increase of unknown ASVs affiliated to *Lactobacillus, Streptococcus, Staphylococcus* and Enterobacteriaceae, as well as *Veillonella dispar* and *Veillonella purvula*. WS_T1 also showed a significant decrease in ASVs affiliated to *Porphyromonas, Paenibacillus, Streptococcus, Actinomyces, Gemellaceae, Granulicatella* and *Abiotrophia*. The microcosms supplemented with sucrose, at timepoint 2 (WS_T2), were characterized by a significant increase in unknown ASVs affiliated to *Staphylococcus Enterococcus, Lactobacillus, Streptococcus Fusobacterium, Atopobium, Neisseriaceae* and *Eikenella*, as well as *Veillonella dispar* and *Streptococcus anginosus*. WS_T2 also displayed a significant decrease of ASVs affiliated to *Bacillus, Porphyromonas* and *Paenibacillus* (Figure 5a and Supplemental Table 6).

We corroborated these biomarkers using indicator species analyses and its indicator value “IndVal” index to detect taxa that specifically characterized treatment (e.g. sucrose supplementation) and each time point (4-way comparisons, Supplemental Table 7). Representative taxa for each studied group showing IndVal index, >0.4, and corroborated using an FDR-adjusted *P*-value (multiple comparisons) are presented in Figure 5b. The rest of the identified indicator taxa (25 in total) and their IndVal and probabilities are listed in Supplemental Table 7. The ASVs with the strongest IndVals were associated with WS_T2, which represented overt lesions; these taxa belonged to *Lactobacillus, Atopobium* and *Enterococcus*. Strong indicator ASVs for NS_T1 include *Abiotrophia* and *Porphyromonas*, while incipient early lesions (WS_T1) corroborated high abundance of *Streptococcus* (Supplemental Table 7).

### Targeted analysis of the central carbon metabolism

LC-MS was used to comprehensively identify and quantify metabolites in the central carbon metabolism – glycolysis pathways (the Embden-Meyerhof-Parnas (EMP) pathway, the pyruvate metabolism, the pentose-phosphate pathway, the Leloir pathway, and the TCA cycle) contained in the biofilms grown on teeth slices for *ex-vivo* caries induction. A total of 18 metabolites were identified and the area under the peak was calculated for each: Acetyl coenzyme A (ACoA), Alpha-ketoglutarate (α-KetoGlu), Citrate, Dihydroxyacetone phosphate (DHAP), Fructose 1,6-bisphosphate (F1,6bP), Fructose 6-phosphate (F6P), Fumarate, Glucose 1-phosphate (G1P), Glyceraldehyde 3-phosphate (G3P), Glucose 6-phosphate (G6P), Galactose-1-phosphate (Gal1P), Lactate, Malate, Phosphoenolpyruvate (PEP), Pyruvate, Ribose 5-phosphate (R5P), Ribulose 5-phosphate (RL5P), Succinate (Succ). For the metabolomic data analysis, the peak area was normalized, log-transformed and pareto-scaled to keep data structure partially intact, non-dimensionless, and closer to the original measurement compared to other scaling methods (59) (Supplemental Figure 6).

The unsupervised Principal Component Analysis (PCA) and supervised Partial Least Squares-Discriminant Analysis (PLS-DA) showed a greater distinction between the treatment groups than between time points (Figure 6a and C respectively). Permutation tests for the PLS-DA model, as calculated by separation distances (1000 permutations), corroborated significant distinction (*p*-value < 0.001) and cross validation for two components showed R2= 0.91 and Q2=0.86 (Supplemental Figure 6). Correspondence between the tested metabolites and samples was depicted in a loadings biplot (Figure 6b) where arrows represent the association of a given metabolite with samples displayed in the PCA ordination (Figure 6a). WS_T2 samples, representing overt lesions, were closely associated with ACoA, DHAB, G3P, G6P, Lactate and Pyruvate, circled in the top dashed blue oval (Figure 6b). Conversely, Fumarate was exclusively associated with samples grown in NS conditions, circled in the bottom left dashed blue oval (Figure 6b). α-KetoGlu, F1,6bP, G1P, PEP, R5P, RL5P were not specifically associated with any tested group (Figure 6 B) and showed the least importance on the PLS-DA projection (Variable Importance in Projection “VIP” scores), which scores metabolites based on their influence in distinguishing each group (Figure 6d, blue oval). Lactate and Pyruvate recorded the highest VIP scores, indicating that they are the most influential metabolites in predicting the phenotypes of interest as shown in the PLS-DA model (Figure 6d). A hierarchically-clustered heat map, with a dendrogram based on Euclidean distances and Average algorithm, showed detailed associations between all tested metabolites and samples, where the samples clustered closely together, first according to the treatments and subsequently by time point (Figure 6e). A heat map focused on average abundances of a metabolite across all samples of a given comparison group collectively summarizes these patterns (Figure 6f). During caries onset, WS_T1, DHAB, G3P, Lactate and Pyruvate were significantly upregulated, which corresponded with a depletion of Fumarate, the only significantly downregulated metabolite compared to the normal NS conditions (Figure 6f).

**Figure 6.**
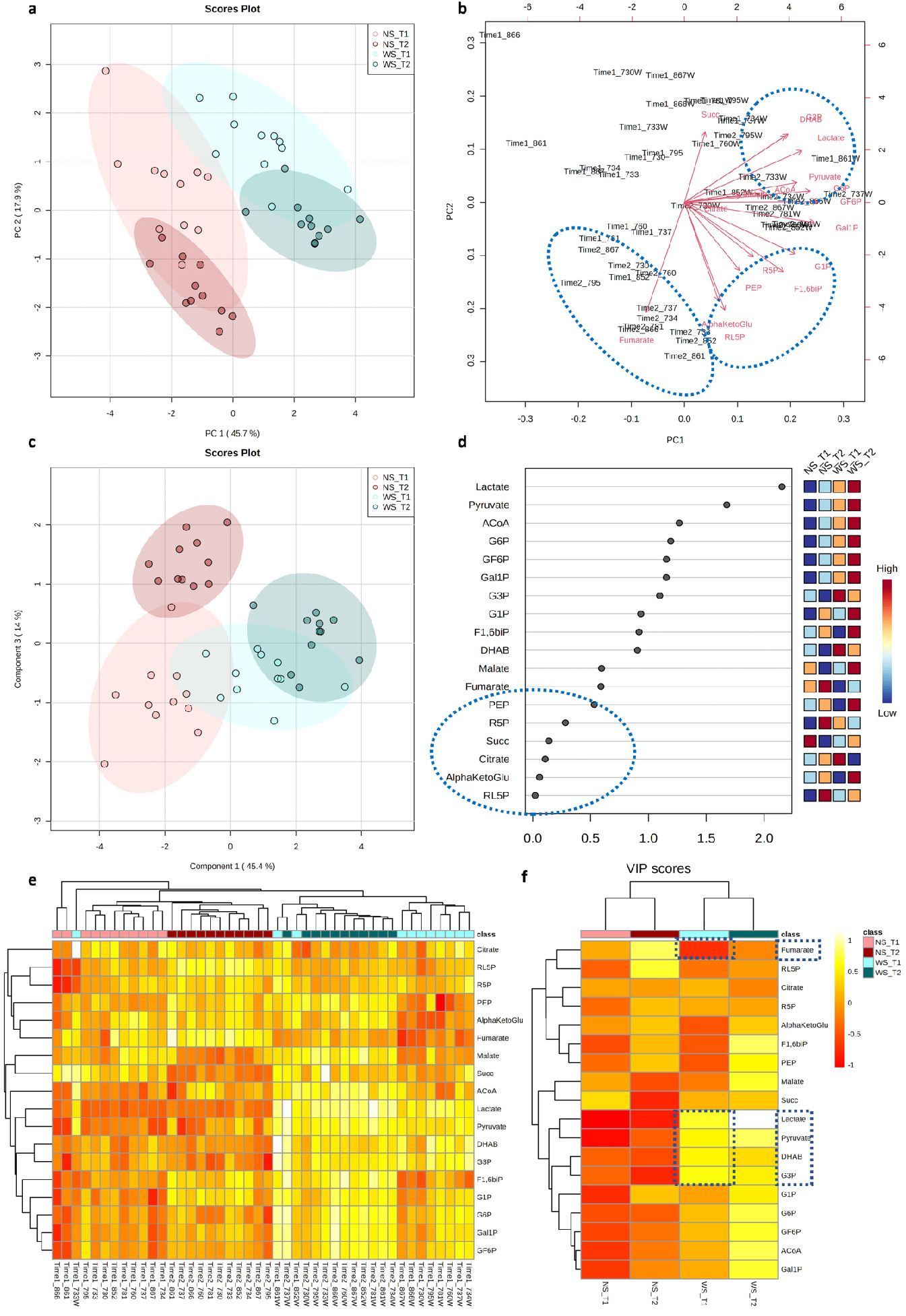
Differential metabolomic profiling of the central carbon metabolism associated with dysbiotic (cariogenic) and non-dysbiotic (control) conditions at the *ex-vivo* dental caries onset and after progression to overt lesions. **A) Principal Component Analysis (PCA)** shows clustering of samples from each group, and color-coded ovals displaying 95% confidence intervals in multivariate space; the red and green shades denote the samples grown in control and dysbiotic conditions respectively, where lighter tones refer to first time points and darker tones refer to the second time points of analysis. The percent of variability accounted for by each axis is indicated. **B) Biplot illustrating the correspondence between metabolites and samples**. Arrows portray the association of specific metabolites with the samples displayed in PCA. The arrow length represents the influence of the metabolite and arrows that have a small angle between them are indicative of metabolites that co-occur with each other. **C) Partial Least Squares-Discriminant Analysis (PLS-DA) for both classification and feature selection**. The permutation and cross-validation tests of the model are detailed in Figure 6–figure supplement 1. **D) Variable Importance in Projection (VIP) scores**. VIP is a weighted sum of squares of the PLS weights, which indicates the importance of each variable or metabolite to the model and to differentiate the groups. VIP values <0.5 show the metabolites that were not influential in this study. **E) Heat map analysis with the dendrogram based on Euclidean distance and Average algorithm**. Columns represent individual tested samples and rows represent 18 targeted metabolites of the central carbon metabolism. The relative abundance of each metabolite is represented by color in each cell. The color-coded groups are presented on the top of the heat map. **F) Average heat map showing differential metabolites per group**. The dashed boxes indicate the upregulated and downregulated metabolites significantly associated with the caries onset. “NS_T1” stands for No Sucrose at Time Point 1-1^st^ phase/caries onset, “NS_T2” stands for No Sucrose at Time Point 2-2^nd^ phase/overt lesions, “WS_T1” stands for With Sucrose at Time Point 1-1^st^ phase/caries onset and “WS_T2” stands for With Sucrose at Time Point 2-2^nd^ phase/over lesions.

We conducted pairwise comparisons of the quantified metabolites in cariogenic and control conditions at caries onset and after progression along the glycolysis pathways (Figure 7). We found 9 metabolites significantly upregulated at the caries onset (ACoA, DHAP, F6P, G1P, Gal1P, G6P, G3P, Lactate, Pyruvate) circled in dashed red ovals (Figure 7). Four of them had key large differences and highly significant *q* values compared to the rest of the metabolites (DHAB “*q*=0.007”, G3P “*q=*“0.005”, Lactate “*q*=7×10^−9^”, Pyruvate “*q*=4×10^−8^”) (Table 1). Fumarate was the only downregulated metabolite and at a significant level at the caries onset (Figure 7) (Table 1). The above four key upregulated metabolites with the caries onset, and the correspondent depletion of fumarate are highlighted in the aforementioned heat maps (Figure 6e and f). For the overt lesions, 11 highly upregulated metabolites and 2 downregulated ones were identified (Table 1). Significance values for the upregulation and downregulation of all metabolites associated with the caries onset and overt lesions compared to collateral normal conditions are listed in Table 1.

**Table 1:** The statistical significance values of the pairwise comparisons of the quantified metabolites in dysbiotic (cariogenic) and non-dysbiotic (control) conditions at each time point. *q* value denotes the *p* value that has been adjusted for the false discover rate (FDR). Double asterisks denote key large differences.

|  | Metabolites/Biomarkers | Caries Onset | Overt Lesions |
| --- | --- | --- | --- |
| 1 | Acetyl-CoA | up regulated ($q=0.01$)* | up regulated ($q=6.95 \times 10^{-4}$)** |
| 2 | Dihydroxyacetone phosphate | up regulated ($q=0.007$)** | up regulated ($q=1.7 \times 10^{-5}$)** |
| 3 | Fructose 6-phosphate | up regulated ($q=0.014$)* | up regulated ($q=1.59 \times 10^{-5}$)** |
| 4 | Fructose 1,6-bisphosphate | | up regulated ($q=2.42 \times 10^{-3}$)** |
| 5 | Fumarate | down regulated ($q=0.015$)* | down regulated ($q=1.59 \times 10^{-5}$)** |
| 6 | Glucose 1-phosphate | up regulated ($q=0.015$)* | |
| 7 | Galactose 1-phosphate | up regulated ($q=0.014$)* | up regulated ($q=9.72 \times 10^{-5}$)** |
| 8 | Glucose 6-phosphate | up regulated ($q=0.014$)* | up regulated ($q=5.05 \times 10^{-5}$)** |
| 9 | Glyceraldehyde 3-phosphate | up regulated ($q=0.005$)** | up regulated ($q=1.59 \times 10^{-5}$)** |
| 10 | Lactate | up regulated ($q=7.18 \times 10^{-9}$)** | up regulated ($q=1.56 \times 10^{-11}$)** |
| 11 | Malate | | up regulated ($q=1.59 \times 10^{-5}$)** |
| 12 | Pyruvate | up regulated ($q=4.26 \times 10^{-8}$)** | up regulated ($q=1.23 \times 10^{-4}$)** |
| 13 | Ribulose 5-phosphate | | down regulated ($q=2.67 \times 10^{-5}$)** |
| 14 | Succinate | | up regulated ($q=1.23 \times 10^{-4}$)** |

**Figure 7.**
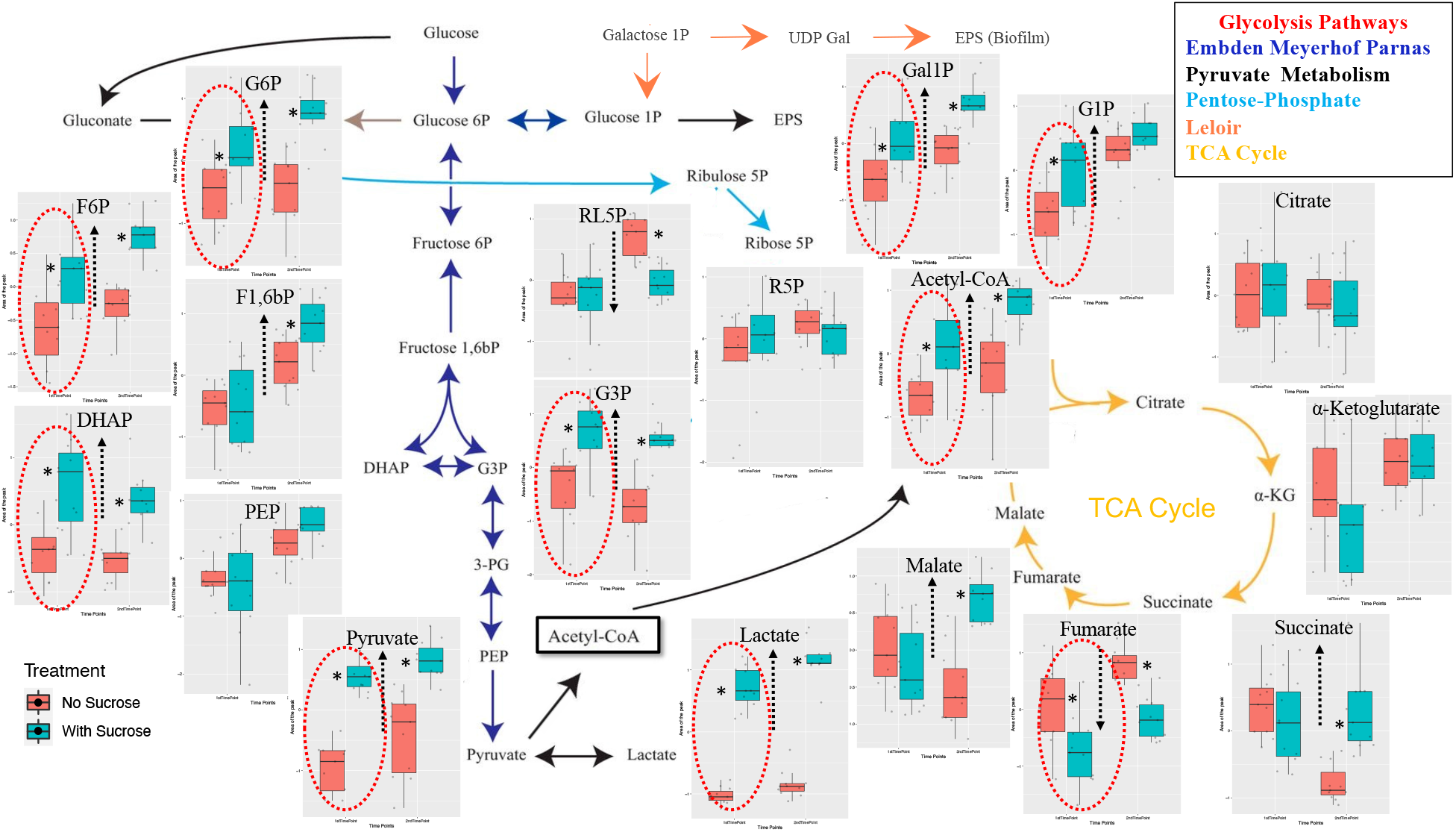
Pairwise comparisons of the profiled and quantified metabolites in dysbiotic (cariogenic) and non-dysbiotic (control) conditions at the *ex-vivo* dental caries onset and after progression along the specified glycolysis pathways. Per each metabolite cluster, the dysbiotic and non-dysbiotic conditions are depicted in green and red shades, respectively, where the first time point-1^st^ phase/caries onset analysis is shown on left and the second time point-2^nd^ phase/overt lesions is on right. The black dashed arrows inside each box plot show the significantly upregulated or down-regulated metabolites at either time point. The red dashed ovals mark the significantly different metabolites, specifically at the first time points that correspond to the caries onset.

Principal coordinates analyses (Bray-Curtis distances) on the targeted metabolome data corroborated differentiation between samples based on the sucrose treatment rather than time elapsed, unlike the temporally-dominating changes observed with the 16S rRNA data (Supplemental Figure 5a). Further, unlike the patterns observed with 16S data, hierarchical clustering of Bray-Curtis distances on the metabolome data showed two distinct groups under health (NS) and cariogenic conditions (WS). This observation indicates the existence of shared metabolic pathways among taxonomically diverse biofilms (Supplemental Figure 5b). The PERMANOVA test for both the metabolome and microbiome profiles showed statical significance for the treatment effect, for the time effect, and for their interaction (Supplemental Figure 5c). However, as expected, the PERMANOVA R^2^ for treatment was higher in the metabolome data compared to 16S data; 0.59 and 0.15 respectively, while the R^2^ for time point was higher in the 16S data compared to metabolome data; 0.24 and 0.10 respectively (Supplemental Figure 5c).

Lastly, we investigated the level of association or correspondence between metabolome and microbiome biomarkers (Figure 8). These associations were initially tested using Procrustes and Mantel tests (Procrustes M^2^ = 0.65, r = 0.59, *p* = 0.001; Mantel r = 0.3, *p* = 0.001, 999 permutations), showing that the two distance matrices (Bray-Curtis) are significantly correlated. Subsequently, we tested associations between specific bacterial taxa (ASVs) and metabolites using *mmvec;* this procedure identifies the highest co-occurrence probabilities; i.e., the metabolites that mostly corresponded with abundance of a given bacterial taxa. Larger positive log conditional probabilities, displayed in red in figure 8, indicate a stronger likelihood of co-occurrence, while low correspondence (negative values), displayed from white to blue, indicate no relationship, but not necessarily a negative correlation (Figure 8). We found strong associations between specific taxa and key upregulated metabolites associated with caries onset, especially Lactate and DHAP. Specifically, strong associations were found between DHAP and *Actinomyces* and between Lactate, DHAP and *Veillonella parvula, Streptococcus, Porphyromonas* and *Granulicatella* (Figure 8). Weaker associations were found between Pyruvate and *Paenibacillus* and *Fusobacterium* (Figure 8). Other strong associations were found between ACoA and *Enterococcus, Actinomyces, Haemophilus parainfluenzae*, and *Paenibacillus*; and between F6P and *Paenibacillus* (Figure 8). No significant associations were found between Fumarate, the only downregulated metabolite in WS-T1and WS-T2, and any of the aforementioned taxa (Figure 8).

**Figure 8.**
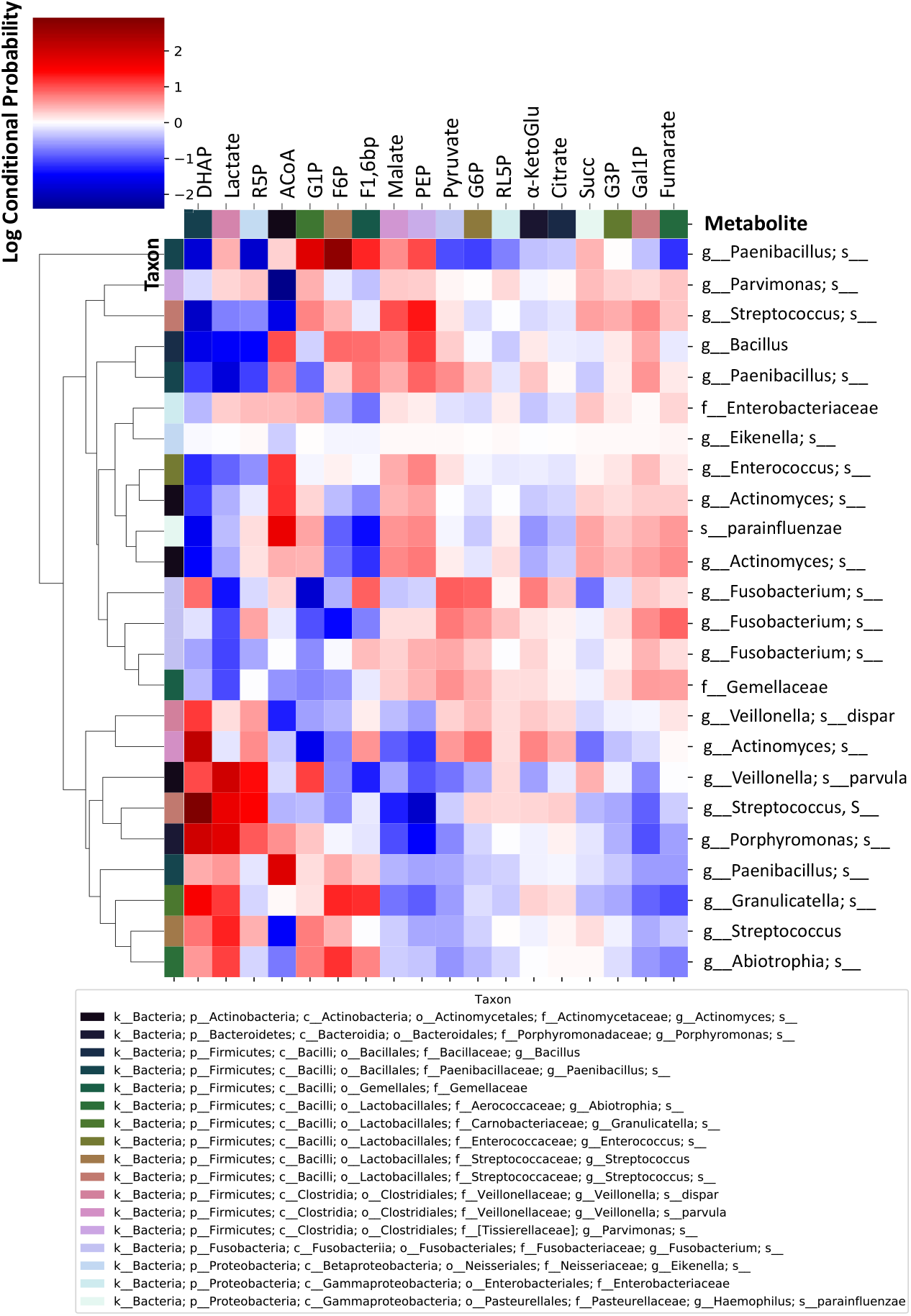
Co-occurrence analyses between microbes and metabolites in supragingival plaque microcosms in dysbiotic (cariogenic) and non-dysbiotic (control) conditions at the *ex-vivo* dental caries onset and after progression. The metabolites are displayed at the top of the heat map while the key taxa (ASVs) and their clustering dendrogram are displayed on the sides. The clustered heat map infers the log conditional probabilities between taxa and metabolites where larger positive conditional probabilities (displayed in red) indicate a stronger likelihood of co-occurrence and low and negative values (displayed from white to blue) indicate no relationship but not necessarily a negative correlation.

## Discussion

Although a dysbiotic state is agreed to be a key factor contributing to the onset of dental caries, our understanding of the functional changes accompanying the transition from a healthy oral ecosystem to the dysbiotic onset of caries is seldomly studied. Given the site-specific and dynamic nature of dental caries, profound understanding requires a longitudinal and multi -level analysis that includes taxonomic assessment (composition) and potential functions (metagenome), and/or active functions (metatranscriptome), and/or encoded functions (metaproteome and metabolome) within the context of this complex oral ecosystem (60) (61). In contrast to metatranscriptome and metaproteome analyses, which represent one aspect of cellular function, metabolomic profiling can provide an instantaneous snapshot of the physiological performance of a given microecosystem. Thus, metabolomic profiling reveals what is actually happening as opposed to the potential for something to happen. Furthermore, the metabolome is the closest link to the phenotype that reflects all the information expressed and modulated by all other Omic-layers along the biological hierarchy (62).

Herein, we present the first integrative downstream analysis of the bacterial compositional and metabolic changes of an *ex-vivo* ecosystem model of supragingival plaque associated with dental caries onset and progression. Our objective was to capture the etiological changes of this disease that are otherwise undetected clinically (Figure 1). We focused on analyzing the compositional and metabolic patterns associated with sucrose-induced pH-changes because of the evident direct relation between organic acids produced by dietary carbohydrate metabolism and the pathogenicity of dental caries (63-65) (Supplemental Table 2).

Our novel *ex-vivo* model allowed us to provide a quantitative analysis of changes in central carbon metabolism captured at the dental caries onset, in tandem with the associated microecological changes. Not only the microcosm model used exhibits a highly diverse community approaching the diversity of human supragingival plaque (26, 27), but also each tested pair of the *ex-vivo* model, the microcosm pairs and teeth substrates pairs, is derived from a single patient. This test-control matching is vital to circumvent the biologically confounding factors inherent to high interindividual variation observed in the oral ecosystem (Figure 1).

Besides, this *ex-vivo* model integrated with a powerful non-invasive advanced bioimaging, MP-SHG, has also enabled us to simultaneously track structural changes in enamel (rich in minerals only), and dentin (comprised of minerals and collagen), aiming to capture the earliest subtle changes taking place in either one. Although dental caries typically starts in the external layer of enamel (66), we tested both aforementioned tissues for two reasons. First, enamel caries could be arrested for years unlike when the lesion reaches the dentin, where it flares up and necessitates quick response through restorative treatments (66). Second, caries can start right in dentin in cases of severe enamel attrition, abrasion and/or erosion as well as in geriatric patients who experienced gingival recession and root surface exposure (67).

The MP-SHG ultrastructural non-invasive imaging was key to set the 1^st^ timepoint that corresponds to the caries onset ; i.e., incipient lesions that could not be otherwise detected (66) (Figure 3 and Supplemental Figure 2). Additionally, MP-SHG enabled the longitudinal study design to track changes in the same samples from the sound start until the detection of overt lesions, going through the critical stage of transition from health to disease (Figures 2, Figure 3, and Supplemental Videos 1). At the caries onset, the major structural disintegration was particularly and consistently observed at the junction between enamel and dentin - DEJ - while the collateral samples derived from same patients and inoculated with the same microcosms, except for the sucrose supplementation, did not display any of these disintegrative signs (Figure 2 and Supplemental Figure 1). As lesions progressed and became overt to visualize, a remarkable band of disintegration was observed with MP-SHG in the dentin side of the DEJ (Supplemental Figure 3 and Supplemental Videos 2). The dentin layer that is just subsequent to the DEJ, 20-30 um, called mantle dentin and typically has more collagen, fewer tubules and less overall mineral than the bulk dentin (68-71). The mantle’s softer structure might explain the earlier effect on this layer compared to other zones of cultured dental substrates. This imaging approach is novel in the field, no references were available to validate the lesions we induced *ex-vivo* compared to clinical carious lesions. Accordingly, we implemented another well-established characterization method in the dental literature, SEM, to validate our *ex-vivo* induced lesions by comparison with well-reported clinical carious lesions at the DEJ area (43, 72-74) (Figure 3 and Supplemental Figure 4). Given that the SEM is an invasive technique and cannot be applied in a longitudinal study design, we implemented it just to characterize each stage of induced lesions individually. Evidently, the SEM micrographs confirmed the carious nature of the induced lesions based on the observed demineralization and disintegration patterns as well as collagen disorganization (Figure 3 and Supplemental Figure 4) (75, 76). Correspondingly, we proceeded to the microbiome and metabolome analyses.

16S rRNA data-driven analyses stratified our tested samples into four clusters (Figure 4). Despite changes in the microbial community composition mainly corresponded to temporal dynamics, with samples tightly clustered along PCo.1 (Figure 4a), both effects of treatment and time were significant as well as their interaction as denoted by PERMANOVA tests. The treatment-based changes were delineated in the log fold-change pairwise comparisons where several ASVs showed consistent and significant changes in relative abundances across the 11 human subject-derived microcosms upon sucrose supplementation (Figure 5a and Supplemental Tables 5 and 6). Some taxa that tolerated the sucrose-induced acidic conditions were significantly abundant at the caries onset (WS_T1), such as *Veillonella dispar, Veillonella purvula*, and unknown ASVs from *Lactobacillus, Enterobacteriaceae, Streptococcus* and *Staphylococcus* (Figure 5a and Supplemental Table 6). However, most of these taxa were also found abundant after the lesions became overt (WS_T2), which makes it challenging to point to specific taxa as responsible or exclusively associated with the disease onset. Our identified taxa are among the most-commonly recognized in previous caries association studies; though, these studies identified taxa from different stages of clinical lesions beyond the onset of the disease (13, 18, 77-79). Other genera that were also highlighted for their strong associations and their important influence in polymicrobial carious lesions are the poorly studied *Fusobacterium and Atopobium* (77, 80-82), which we also identified but in the overt lesion stage (Figure 5a and Supplemental Table 6).

Furthermore, indicator taxa analyses (IndVal > 0.6) showed only a few ASVs faithfully distinguishing NS_T1 and in WS_T2, which correspond to early controls and overt lesions (Figure 5b and Supplemental Table 7). Only ASVs affiliated to the *Streptococcus* genus were associated with caries onset (WS_T1), but exact species could not be identified; noting a significant limitation of the 16S rRNA sequencing. Nonetheless, even if the sequencing is capable of providing species-level resolution, the presence of one or more species would not ascertain the causative organisms for the disease; first, because bacterial abundance is not indicative of their activity (17, 77, 83); and second, because the heterogenic polymicrobial nature of this disease significantly varies among individuals (7). Given these two requirements, the indicator ASVs reported herein may be co-incidentally shared among the tested subjects, but they may not give reliable inference about their actual activity within the tested ecosystem.

For functional profiling, we looked at discernable signatures in the central carbon metabolism at health and cariogenic conditions, while testing their associations with the microbiome taxonomic profiles. This approach previously revealed that stable metabolic pathways may exist despite taxonomic heterogeneity across individuals for supragingival plaque (22, 84).

Interestingly, we found that 6 metabolites - ACoA, DHAB, G3P, G6P, Lactate and Pyruvate - specifically associated with samples incubated in cariogenic conditions, whereas one metabolite – Fumarate - was exclusively associated with the controls (Figure 6b). Our findings align to a great extent with previous *in-vivo* studies for metabolome profiling of the supragingival dental plaque before and after glucose supplementation (65, 85). Obviously, the *in-vivo* environment is more taxonomically rich than the microcosm ecosystem; however, these findings support that oral microbial communities share conserved specific metabolic pathways under dysbiotic conditions, despite heterogenous taxonomic composition (65, 85, 86). Additionally, these findings further validate our microcosm model, which previously showed conserved proteome patterns despite taxonomic variability (23).

Worthy of mention, not all identified key metabolites showed the same influence in discriminating between conditions. Lactate was found to be the most powerful differentiating metabolite followed by Pyruvate (Figure 6d). The rest of the studied 18 metabolites showed a gradually decreasing influence, where 6 of them showed a minimal influence; (Figure 6d) and were also found to be not specifically associated with any tested group (Figure 6b).

For the caries onset in particular, five significantly expressed metabolites were observed differentiating caries onset from the control condition, and progression to overt lesions: Lactate, Pyruvate, DHAB, G3P were upregulated, while Fumarate was downregulated (Figure 6e and f and Table 1). Not only the aforementioned 5 key metabolites were significantly different at the caries onset, but also 5 more metabolites - ACoA, F6P, G1P, Gal1P, G6P - were found significantly upregulated at the same time point (WS_T1) (Figure 7 and Table 1). However, these additionally upregulated metabolites did not show high statistical significance as the 5 key metabolites featured in the heat maps (Figure 6e and f and Table 1), nor were they among the metabolites with the highest VIP scores (Figure 6d). As the lesions progressed (WS_T2), a list of 13 metabolites showed high statistical significance; 11 upregulated and 2 downregulated (Figure 7 and Table 1). These finding suggest that the high abundance of Lactate, Pyruvate, G3P, and DHAB, along with the depletion of Fumarate, are enough to denote the onset of dysbiosis and potential incipient lesions existence.

These data are crucial to devise timely interventions with offsetting clinical strategies; such as controlling dietary and hygienic habits (87) and/or developing modern therapeutics for caries prevention or arresting before the disease progresses to overt lesions, which would necessitate invasive restorative dental treatments. Some contemporary approaches include probiotics that antagonize acidogenic/aciduric species (88), targeted antimicrobials that suppress specific pathogens (89), modulating the early acquired enamel pellicle that governs the succession of biofilms (90), and developing alkali production therapeutics, such as arginine, to neutralize glycolytic acids (91).

The upregulation of some metabolites along the glycolysis pathways is self-explanatory based on the fermentation of carbohydrates induced by bacterial metabolism. However, the depletion of Fumarate was an interesting finding that triggered our curiosity (Figure 7 and Table 1). Fischbach and Sonnenburg systematically explained this phenomenon in the context of how anaerobic bacteria generate energy (ATP), maintain redox balance, and acquire carbon and nitrogen to synthesize primary metabolites (92). They elucidated how Fumarate is key for anaerobic ATP synthesis in the final step of the primitive electron transport chain through its reduction to succinate, pointing to this metabolite as the most common terminal electron acceptor for anaerobic respiration (93). Since biofilms were grown in an aerobic environment, as it happens with supragingival plaque *in-vivo*, excessive Fumarate consumption could be attributed to the presence of some strict anaerobic species within the microbial community - such as *Veillonella* (in WS_T1 and T2) and *Fusobacterium* (in WS_T2) *-* which strive to maintain their survival and energy production as aforementioned. Intriguingly, Ribulose-5-phosphate, also showed significant depletion at a later stage, when the lesions became overt (Figure 7 and Table 1). The mechanisms behind depletion of this metabolite are unclear; however, ribulose-1,5-bisphosphate - the product of the phosphorylation of ribulose-5-phosphate- has been found to be the most important CO_2_ fixing pathway in prokaryotes, particularly around oxic/anoxic (free oxygen containing/free oxygen lacking) interfaces (94) that develop as a consequence of oxygen consumption (95). Collectively, Ribulose-5-phosphate consumption seems to be also involved in bacterial adaptation mechanisms used for managing CO_2_ deficiency at an advanced stage of biofilm maturations. We sought to shed light on microbe-metabolite associations behind the metabolic profiles observed (Figure 8). For example, we found strong co-occurrence patterns between Lactate and DHAB, two of the 5 key metabolites associated with the caries onset, and *Veillonella parvula, Streptococcus, Porphyromonas* and *Granulicatella*. Interestingly, although the association of *Streptococcus* and *Veillonella parvula* with the caries onset were recapitulated in the IndVal and Volcano plot analyses, respectively, neither *Porphyromonas* nor *Granulicatella* were found to characterize cariogenesis. Another example is the strong co-occurrence detected between DHAB and an *Actinomyces* ASV (Figure 8), where the latter was not significantly associated with cariogenic conditions at any time point (Figure 5a and Supplemental Tables 5 and 6). Furthermore, Pyruvate, another strong caries onset biomarker, was found associated with *Fusobacterium* (Figure 8) which was identified demarcating the overt stage taxonomy (Figure 5a and Supplemental Tables 5 and 6).

These findings further imply that the mere abundance and/or presence of specific bacteria does not reflect their active role in disease initiation or progression, while the functional profiles/outcomes could be central in controlling the course of the disease (17, 77, 83, 96). Moreover, our findings confirm that taxonomically similar microbiomes may have different metabolic roles in the supragingival microenvironment (97). This scenario is also supported by observations that two ASVs from the same genus; *Actinomyces*, show different patterns of association with DHAB (Figure 8). Likewise three different *Streptococcus* ASVs show various co-occurrence probabilities with Lactate, spanning from no-relationship to strong co-occurrence (Figure 8). In addition, given the limitations of short amplicon sequencing approaches to resolve strain identity, the data may also show that the role of oral bacteria in caries onset and progression is also characterized by fine-level strain or variant dynamics within specific bacterial taxa (21).

Still, all the aforementioned examples of the bacteria associated with the key metabolites are acidogenic/aciduric and have evident roles in dental caries course as reviewed in (78). As such, acidogenic, and acid tolerant bacteria are more likely to contribute to the caries process than other microbiome residents. Nonetheless, as shown here, acidogenic roles can be carried out by different species in different individuals and populations (78).

### Study Limitations

The use of 16S rRNA sequencing techniques poses limitation due to its inability to reveal strain-level characterization of bacterial taxa associated with carries onset or progression (98). Although we identified several ASVs affiliated to same genera, such as *Streptococcus, Actinomyces*, and *Fusobacterium*, displaying different co-occurrence patterns with the metabolomic biomarkers, their precise taxonomic identification and associated metabolic roles in the studied microenvironment could not be resolved (97). For example, *Streptococcus* comprises both commensals, like *S. gordonii* or *S. sanguinis*, and acidogenic and/or aciduric strains, like *S. mutans* or *S. sobrinus* (78, 89), which we could not determine using 16S rRNA short amplicon sequencing. Alongside, the *ex-vivo* model cannot be taken as a surrogate of actual *in-vivo* conditions, especially when considering true microbial diversity. This limitation, in turn, could have influenced the metabolite pool detected in the system. Regardless, we believe the polyphasic approach used, in tandem with the controlled conditions brought by the carefully validated *ex-vivo* model, allowed us to present functional biomarkers, free from the lifestyle confounders imposed by *in-vivo* settings, of the dental caries onset that is otherwise undetected clinically.

## Conclusions

In this study, we implemented a novel longitudinal *ex-vivo* model with the aid of advanced non-invasive and label free MP-SHG bioimaging to induce dental caries experimentally, and characterize the very early signs of the disease, which correspond to undiagnosable subclinical lesions. We analyzed the microbial communities at the disease onset and after progression, using 16S rRNA short amplicon sequencing and central carbon metabolomics. This combined approach allowed us to simultaneously characterize microbiome changes and functional phenotypes associated with the disease, while elucidating associations between microbial biomarkers and metabolic outcomes. Our data revealed five key metabolites significantly expressed with the induced caries onset that did not necessarily co-occur with the most abundant taxa identified under the same condition. This study confirms the crucial role of bacterial activity over their taxonomic abundance in controlling caries pathogenesis, aligning with previous findings using other functional omics platforms. The biomarkers we report for the onset of caries can be key to prevent/arrest the disease at its early stage before it progresses to overt lesions, so that invasive restorative treatments as well as massive expenses in healthcare budgets can be prevented.

## Contributors

- Conceptualization: DM, TM, JR, CA, AG
- Data curation: DM, AS, BW, JP
- Formal analysis: DM, AS, JP, CA, AG
- Funding acquisition: DM, CA, AG
- Investigation: DM, AS, TM, JP, CA, AG
- Writing-original draft: DM
- Writing-review and editing: all authors

## Acknowledgements

The authors acknowledge Julie Kirihara from the University of Minnesota Spectrometry and Proteomics Center for helping in optimizing the metabolites extraction protocols specifically customized for central carbon metabolism byproducts. The authors thank Kayla Law, Kylene Guse, and Samuel Davison from Gomez lab for helping in processing the 16S rRNA row data, MetaboAnalyst software orientation and technical support for the DNA samples tracking and submission to the genomics center, respectively. MP-SHG imaging was performed at the University of Minnesota Imaging Centers (http://uic.umn.edu) with the assistance of Jason Mitchell. The authors thank the labs of Dr. Mansky, Dr. Herzberg, and Dr. Jensen for their lab resources they provide for the DNA and metabolites extraction as well as Dr. Carrera and Mrs. Chen from Dr. Rudney’s lab for helping in collecting the molars and managing the supragingival dental plaque microcosms. This research study was supported by the National Institute for Dental and Craniofacial Research of the National Institutes of Health to Dr. Moussa (R90-DE023058) through the Minnesota Craniofacial Research Training (MinnCResT) Program, Dr. Aparicio (R01-DE026117), and Dr. Gomez with funds from the College of Food Agriculture and Natural resource Science, at the University of Minnesota. The funding bodies had no role in study design, analysis, and interpretation of data; in the writing of the report; and in the decision to submit the article for publication. The content is solely the responsibility of the authors and does not necessarily represent the official views of the National Institutes of Health.

## Data sharing statement

All data needed to evaluate the conclusions in the paper are present in the paper and/or the Supplementary Materials.

