## SupplementaryMaterial for "Functional Biomarkers *of Ex-vivo* Dental Caries Onset"

### Supplemental Figures

| Tooth Pair No.# | Control<br>DEJ Area<br>(Minerals-Enamel/Collagen-Dentin) | | Caries Onset Induction<br>DEJ Area<br>(Minerals-Enamel/Collagen-Dentin) | |
|---|---|---|---|---|
| | Pre-inoculation<br>Sound Tooth Slice | 1st Time Point (88h)<br>No Sucrose (NS) | Pre-inoculation<br>Sound Tooth Slice | 1st Time Point (88h)<br>With Sucrose (WS) |
| #1 | 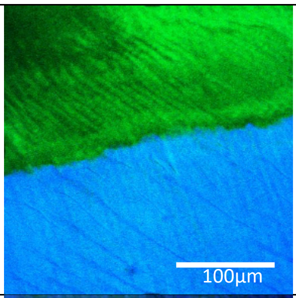  | 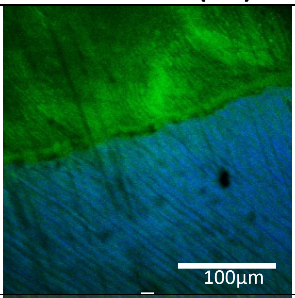  | 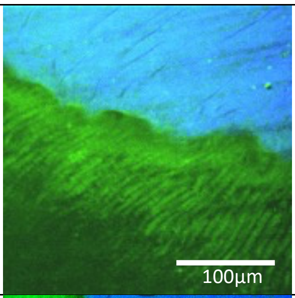  | 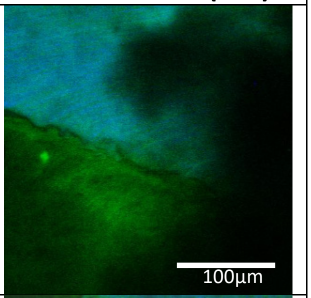  |
| #2 | 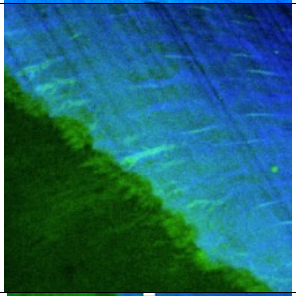 | 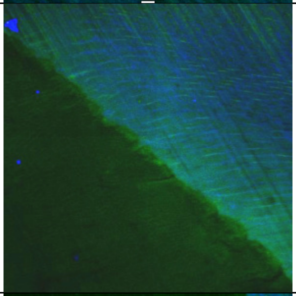 | 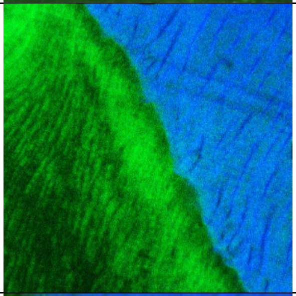 | 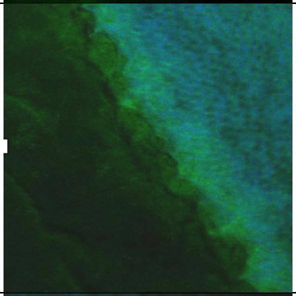 |
| #3 | 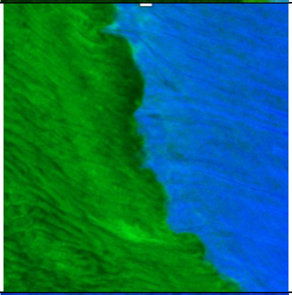 | 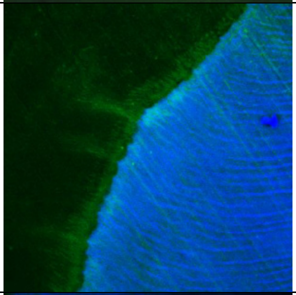 | 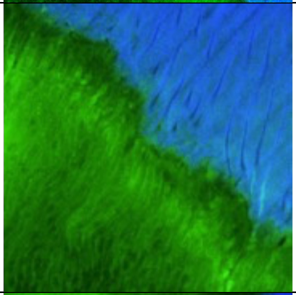 | 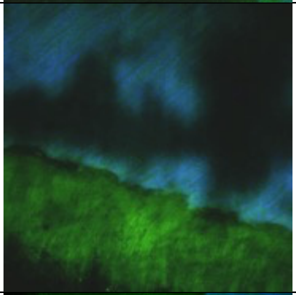 |
| #4 | 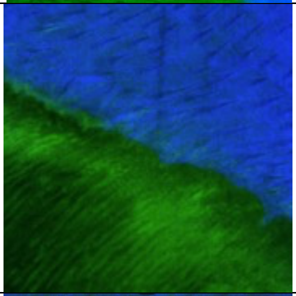 | 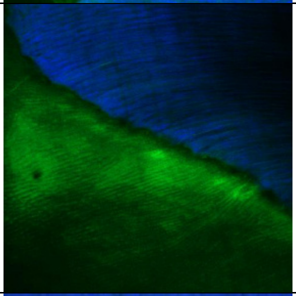 | 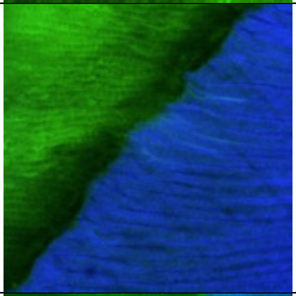 | 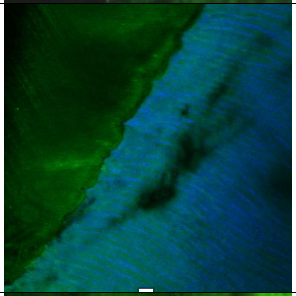 |
| #5 | 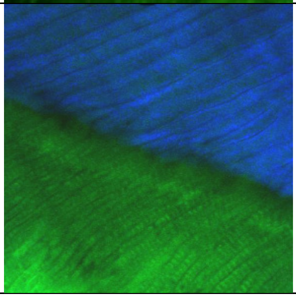 | 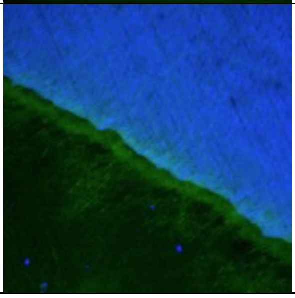 | 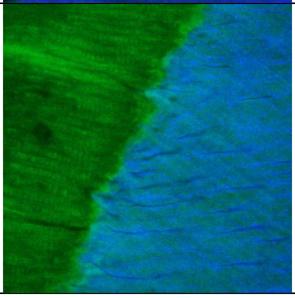 | 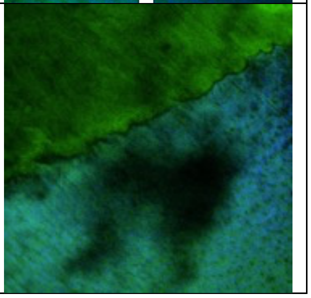 |

| | | | | |
|---|---|---|---|---|
| #6 | 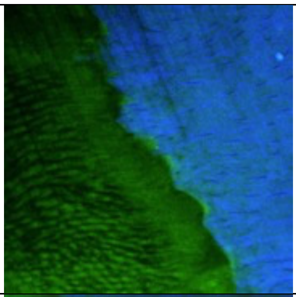 | 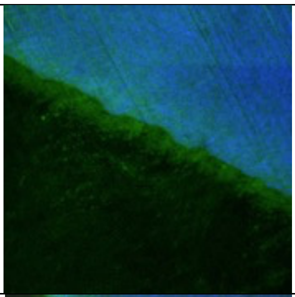 | 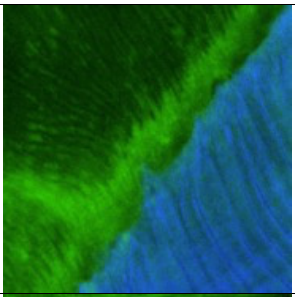 | 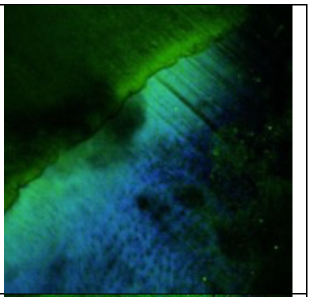 |
| #7 | 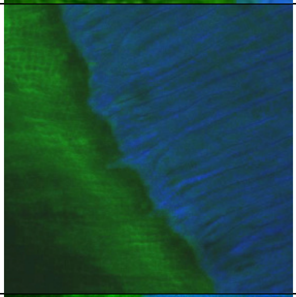 | 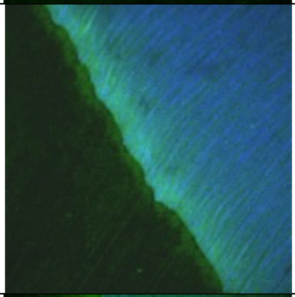 | 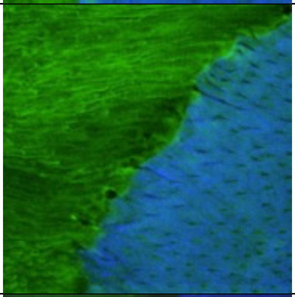 | 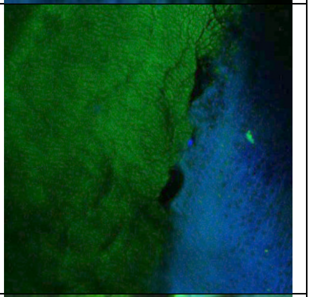 |
| #8 | 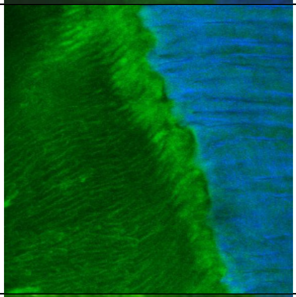 | 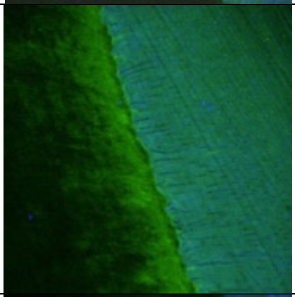 | 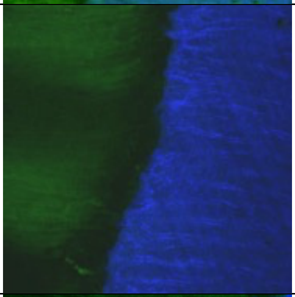 | 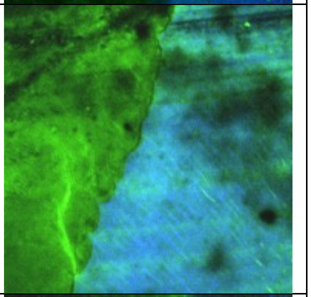 |
| #9 | 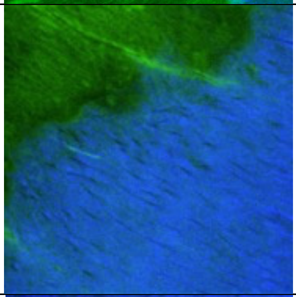 | 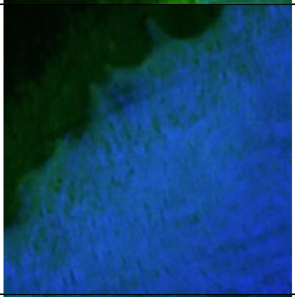 | 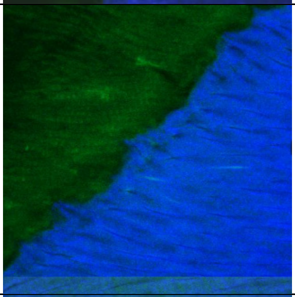 | 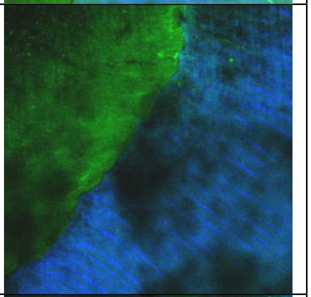 |
| #10 | 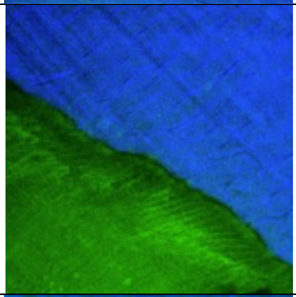 | 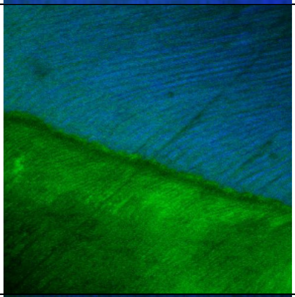 | 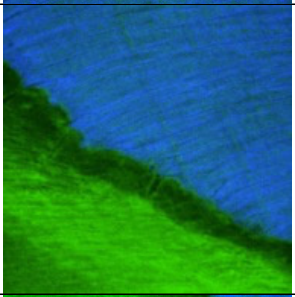 | 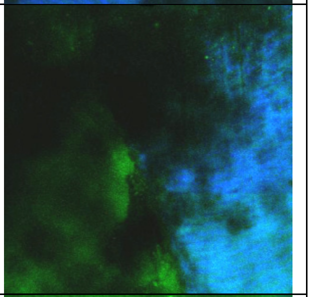 |
| #11 | 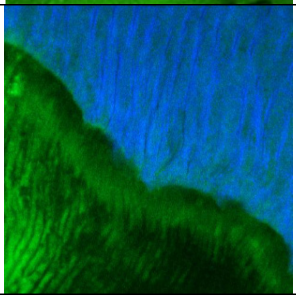 | 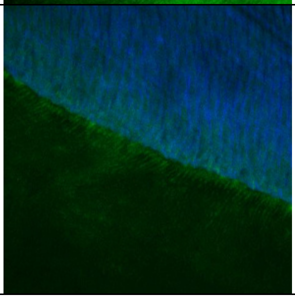 | 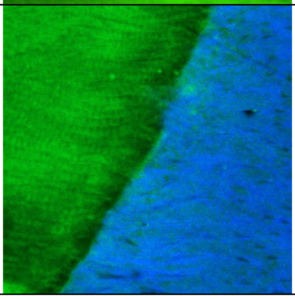 | 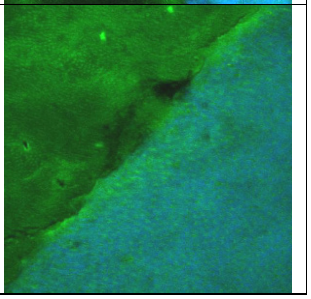 |

**Supplemental Figure 1: Multiphoton-second harmonic generation (MP-SHG) examination of the dentin-enamel junction (DEJ) area of all tested samples before and at the 1$^{st}$ time point of analysis (88h) in control (No Sucrose "NS") and dysbiotic condition (With Sucrose "WS") that corresponds to the dental caries onset.** Each pair of teeth specimens was sliced from a single tooth and inoculated with a pair of supragingival plaque microcosm, without or with sucrose, which originated from one single patient. The green signals show the MP autofluorescence emitted from the mineralized phase (mainly enamel). The blue signals show the SHG exclusively emitted from the collagen network of dentin.

| Tooth Pair No.# | Pre-culturing | | First Time Point (88 hours) | | Second Time Point (17 days) | |
|---|---|---|---|---|---|---|
| | No Sucrose (NS) | With Sucrose (WS) | No Sucrose (NS) | With Sucrose (WS) Caries Onset | No Sucrose (NS) | With Sucrose (WS) Overt Lesions |
| #1 | 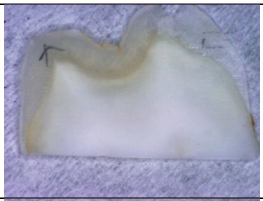 | 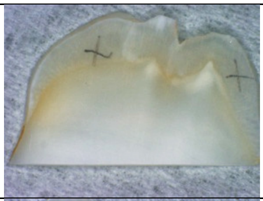 | 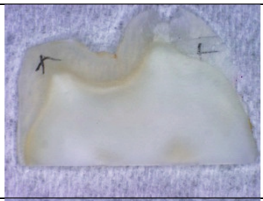 | 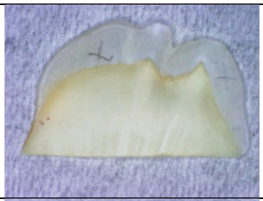 | 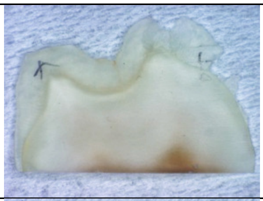 | 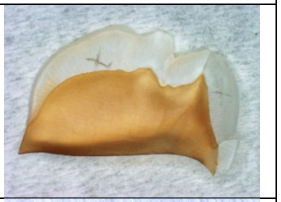 |
| #2 | 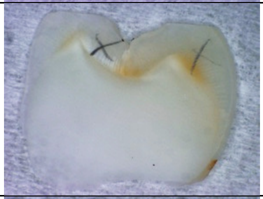 | 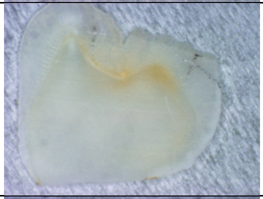 | 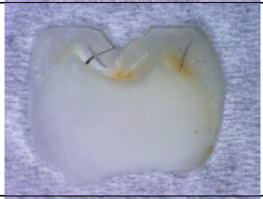 | 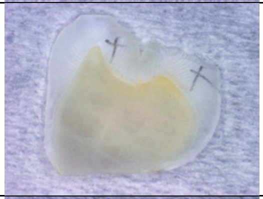 | 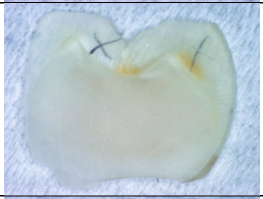 | 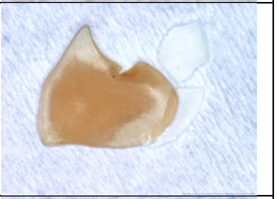 |
| #3 | 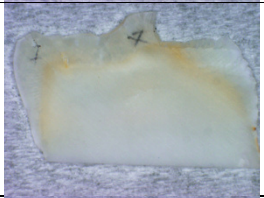 | 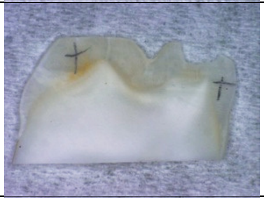 | 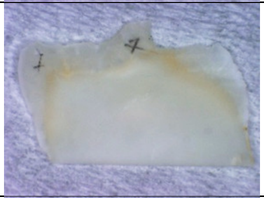 | 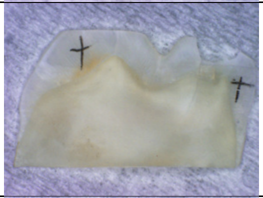 | 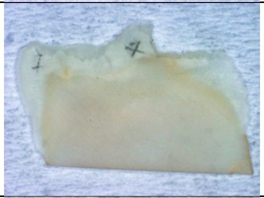 | 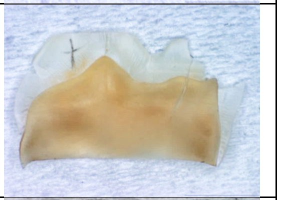 |
| #4 | 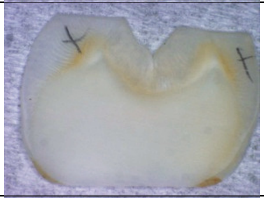 | 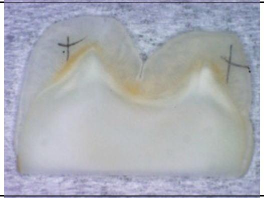 | 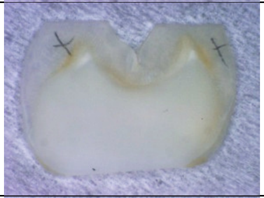 | 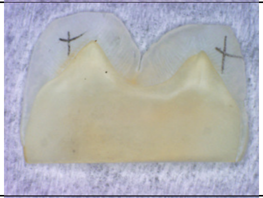 | 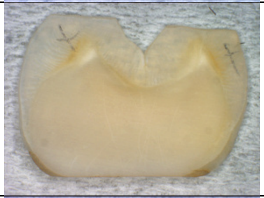 | 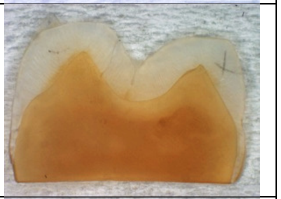 |
| #5 | 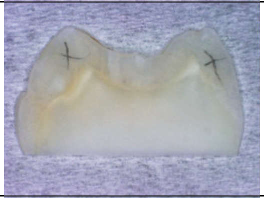 | 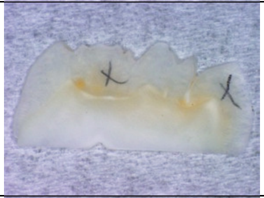 | 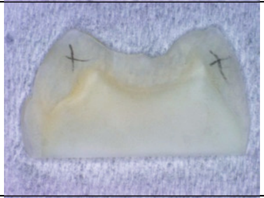 | 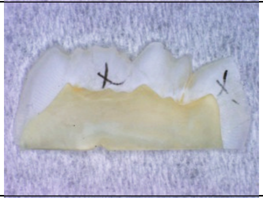 | 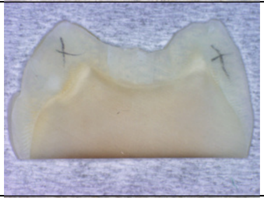 | 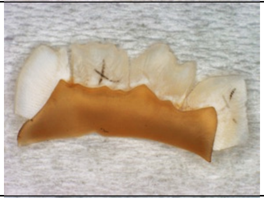 |
| #6 | 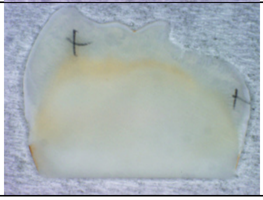 | 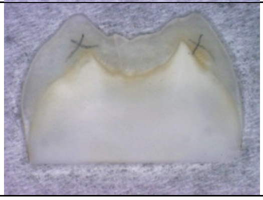 | 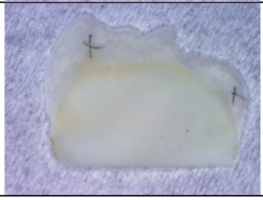 | 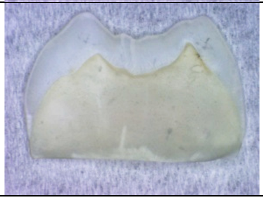 | 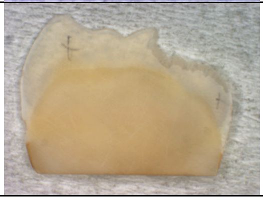 | 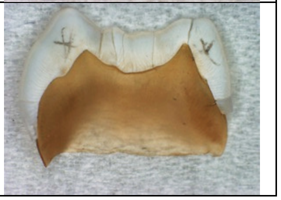 |

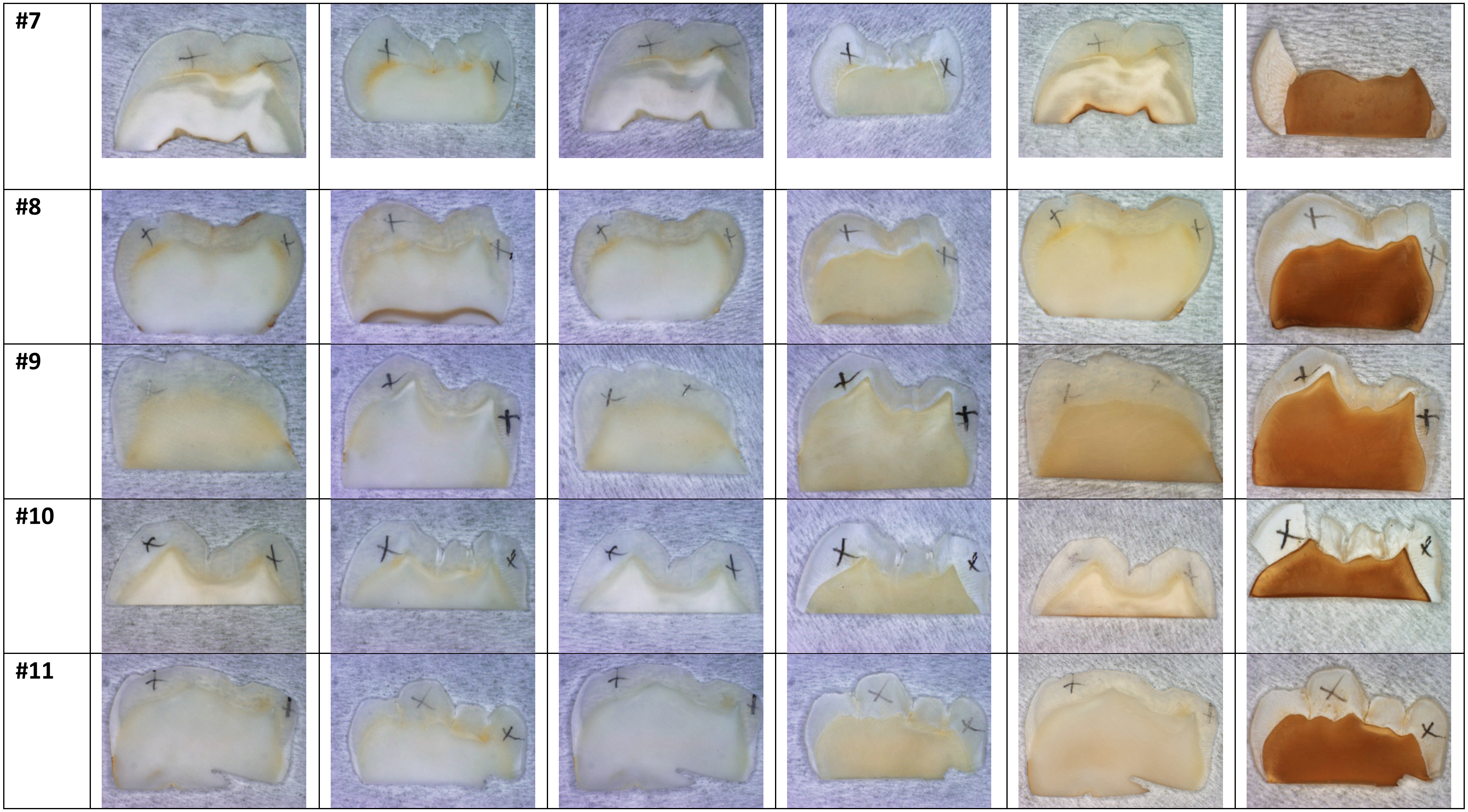


**Supplemental Figure 2: Stereomicroscope screening of all tested teeth specimens before and along the *ex-vivo* caries induction course at each time point of analysis in control (No Sucrose) and dysbiotic (With Sucrose) conditions.** Each pair of teeth specimens was sliced from a single tooth and inoculated with a pair of supragingival plaque microcosm, without (NS) or with sucrose (WS), that originated from a single patient. The NS specimens did not show any signs of developed caries lesion at any time point, as shown on the left column for each time point. In WS specimens, no caries lesions could be distinctly visualized after 88 hours, corresponding to the caries onset; as shown on the right column for the first time point. Distinct signs of caries lesions (discoloration, demineralization, softening, decomposition and cracking) were visualized in all WS samples after 17 days of the caries induction course.

**MP-SHG Monitoring of the Induced Caries Overt Lesions**
**DEJ Area (Minerals-Enamel/Collagen-Dentin)**

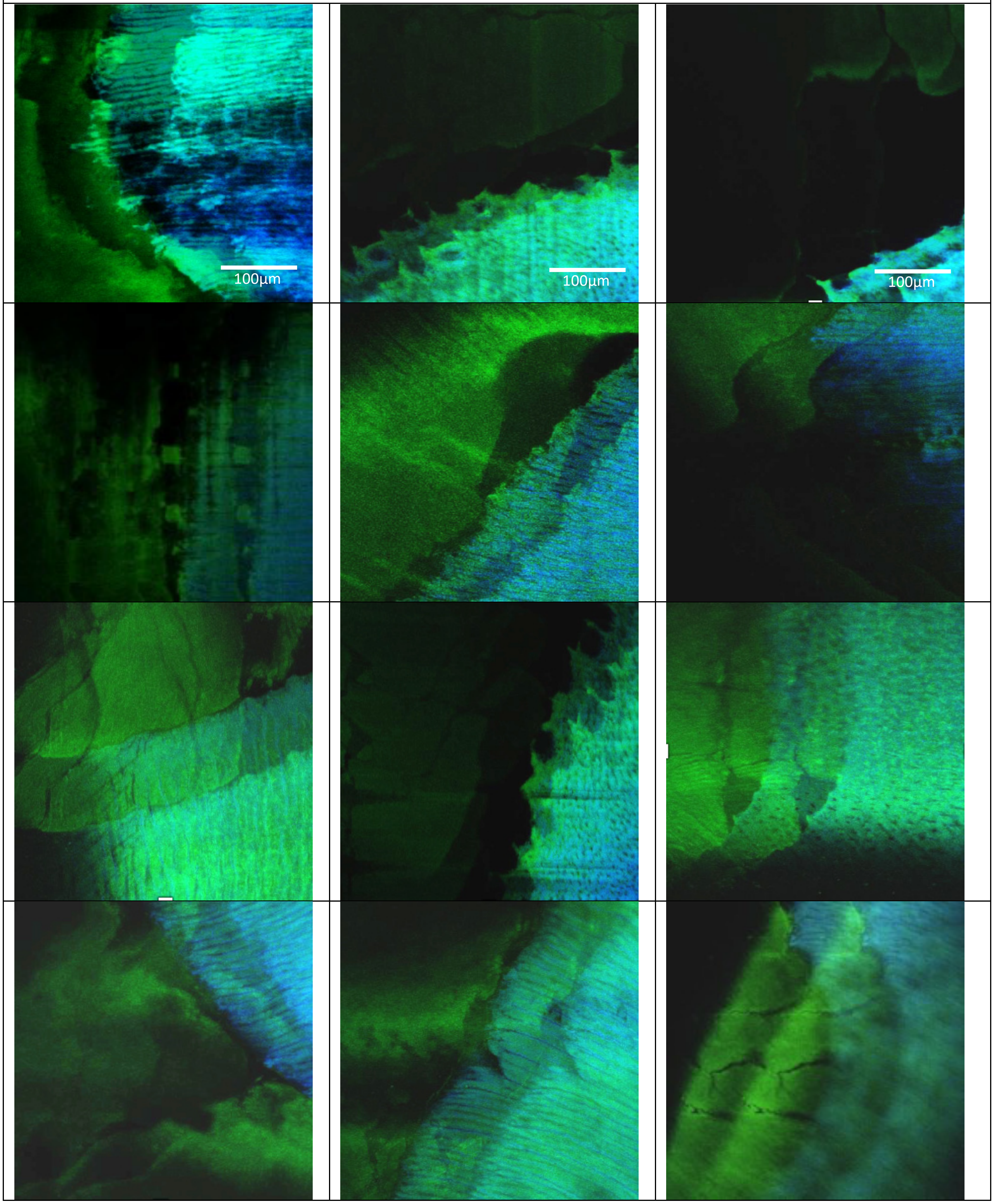


**Supplemental Figure 3: Multiphoton-second harmonic generation (MP-SHG) examination of the dentin-enamel junction (DEJ) area of all tested samples after the *ex-vivo* induction of overt caries lesion (17 days).** Representative Z-stacks of induced overt lesions are displayed in Figure 2-video supplement 2. The green signals show the MP autofluorescence emitted from the mineralized phase (mainly enamel). The blue signals show the SHG exclusively emitted from the collagen network of dentin.

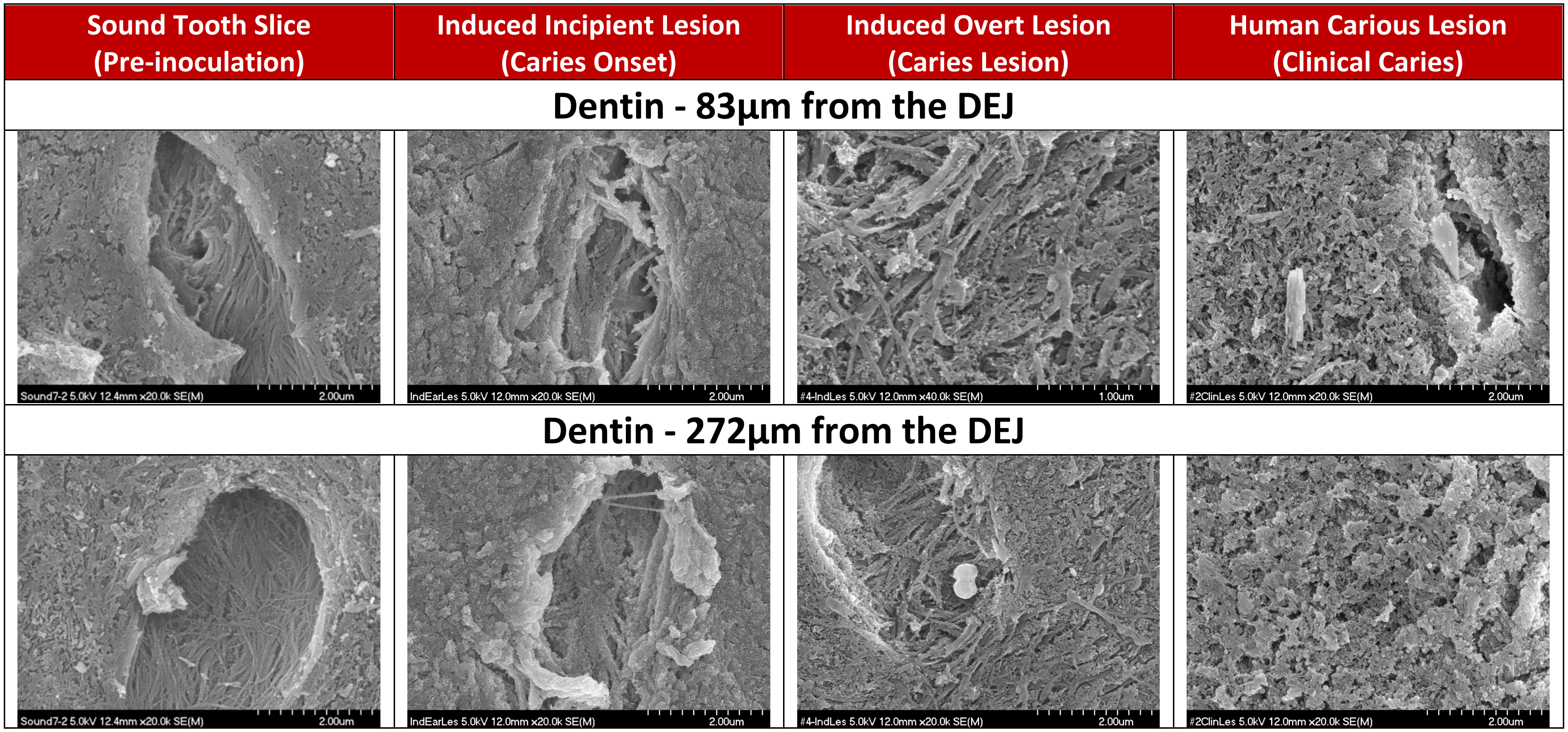


**Supplemental Figure 4: SEM ultrastructural characterization of the stages of induced ex-vivo caries lesion in comparison to the clinical caries lesion at further distances form the dentin-enamel junction (DEJ).** The micrographs showed the gradual degradation of the peritubular dentin with areas of discontinuation along the perimeter of the tubules accompanied with collagen fibers disorganization along the cariogenesis course and at further distances from the DEJ (at 83 µm in the top row and at 272 µm in the bottom row). The induced overt lesions showed less degenerative changes compared to clinical caries at further distances from the DEJ (272 µm) compared to DEJ-closer lesions portrayed in the top row (83 µm from the DEJ) and in Figure 3 (just beneath the DEJ).

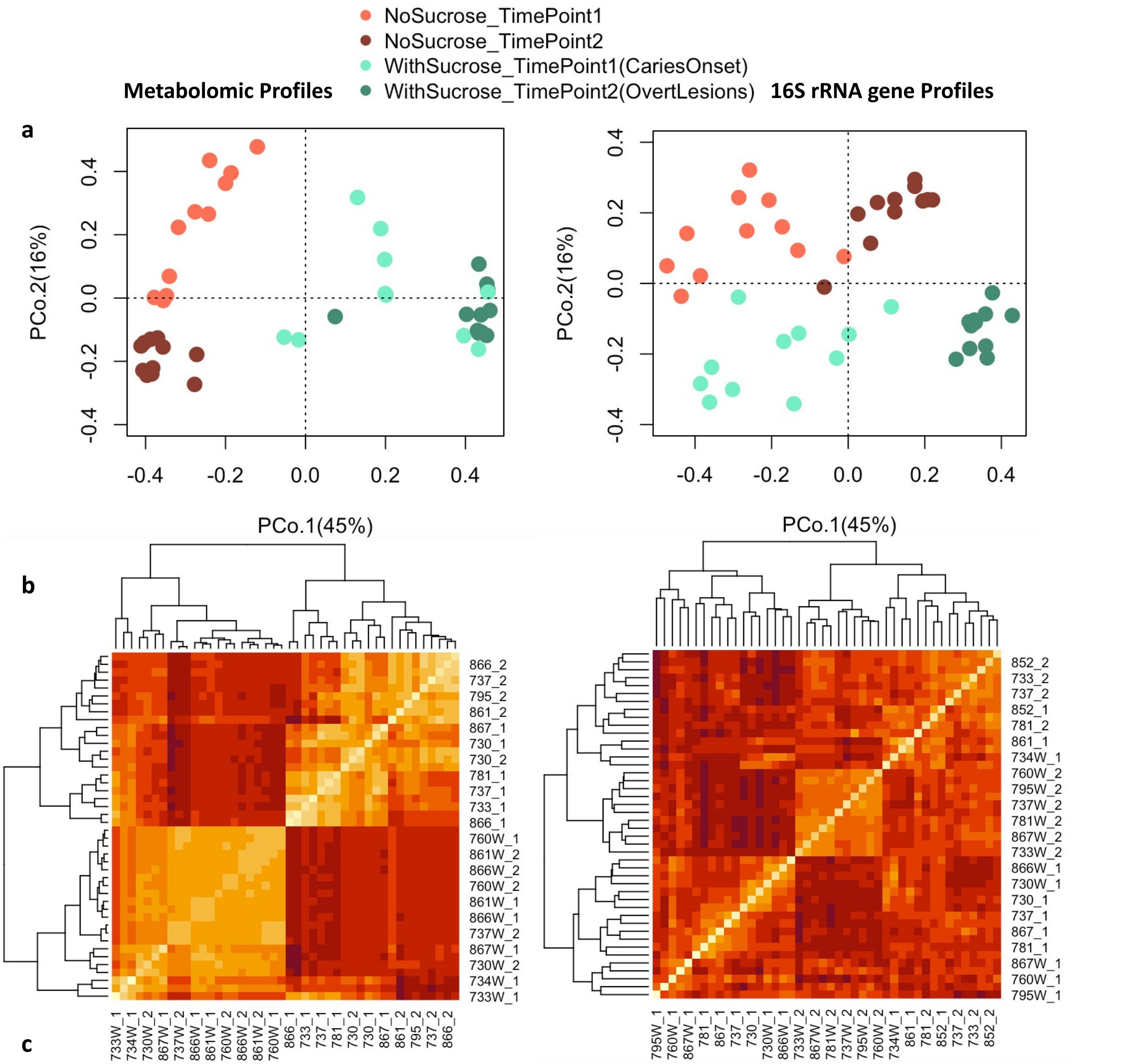


c

| | Df | Sums Of Sqs | Mean Sqs | F.Model | $R^2$ | Pr(>F) |
|---|---|---|---|---|---|---|
| **Treatment** | 1 | 4.11 | 4.11 | 101.18 | 0.59 | 0.001 *** |
| **Time Points** | 1 | 0.73 | 0.73 | 17.98 | 0.10 | 0.001 *** |
| **Treatments: Time Points** | 1 | 0.54 | 0.54 | 13.26 | 0.077 | 0.001 *** |
| **Residuals** | 40 | 1.63 | 0.04 | | 0.23 | |
| **Totals** | 43 | 7.01 | | | 1.00 | |

| | Df | Sums Of Sqs | Mean Sqs | F.Model | $R^2$ | Pr(>F) |
|---|---|---|---|---|---|---|
| **Treatment** | 1 | 1.59 | 1.59 | 10.83 | 0.15 | 0.001 *** |
| **Time Points** | 1 | 2.48 | 2.48 | 16.87 | 0.24 | 0.001 *** |
| **Treatments: Time Points** | 1 | 0.398 | 0.398 | 2.71 | 0.038 | 0.012 * |
| **Residuals** | 40 | 5.87 | 0.15 | | 0.57 | |
| **Totals** | 43 | 10.34 | | | 1.00 | |

**Supplemental Figure 5: Heterogeneity of supragingival plaque microcosms metabolome (left panels) and microbiome (right panels) profiles in dysbiotic (cariogenic) and non-dysbiotic (control) conditions at the *ex-vivo* dental caries onset and after progression**. **a)** Principal Coordinate analysis (PCoA) of the targeted central carbon metabolites and 16S rRNA sequencing reads based on relative abundance. **b)** Clustering co-occurrence plots of the metabolome and microbiome samples ordered along the axes according to the co-occurrence matrix. The more similar the sample profiles, the closer they are together on the axis and the lighter the color on the heatmap. **c)** Permutational multivariate analysis of variance (PERMANOVA) statistical test for the metabolome and microbiome profiles showing the exclusive effect of the induced dysbiosis, the exclusive effect of the time, and their interaction.

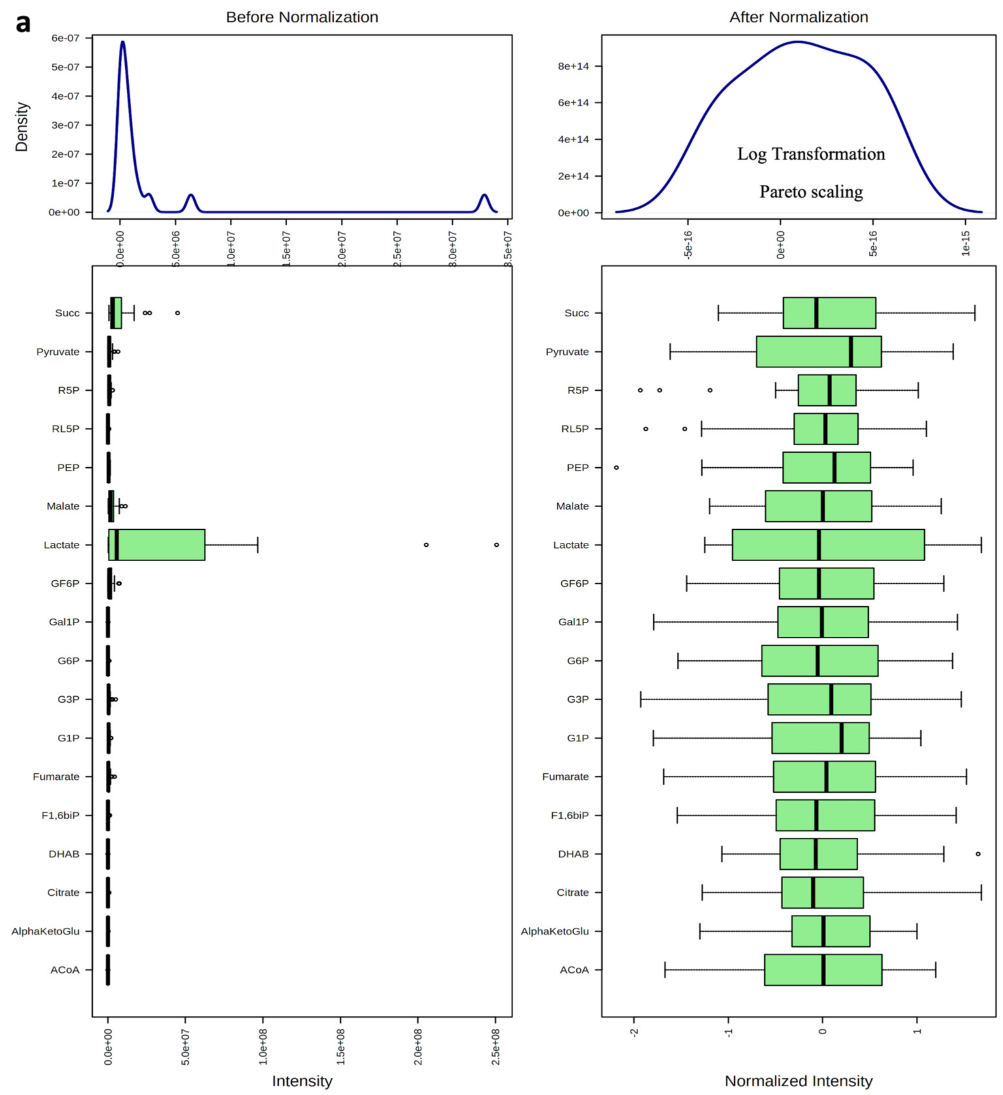


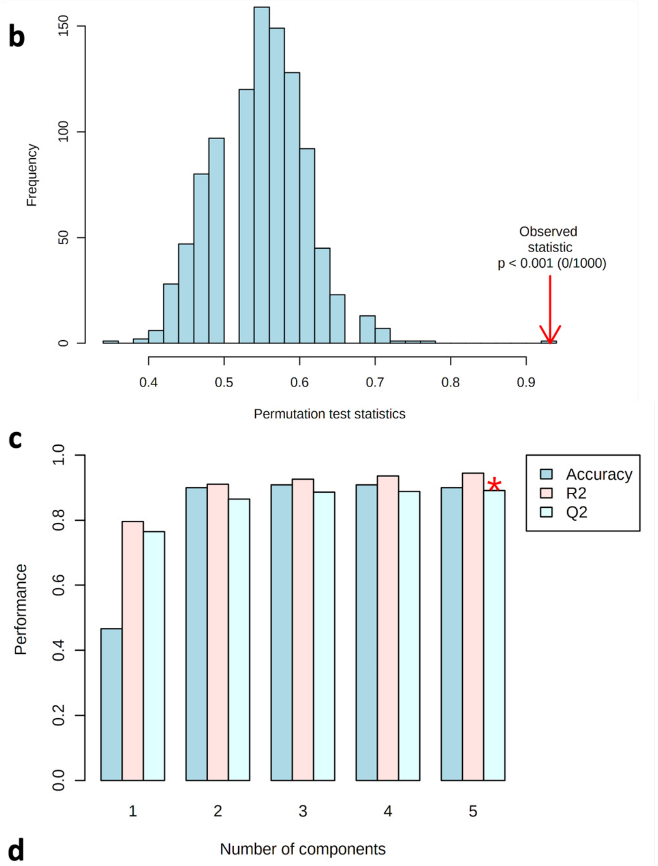


d

| Measure | 1 comps | 2 comps | 3 comps | 4 comps | 5 comps |
|---|---|---|---|---|---|
| Accuracy | 0.46667 | 0.9 | 0.90833 | 0.90833 | 0.9 |
| R2 | 0.79549 | 0.91017 | 0.92566 | 0.93533 | 0.94486 |
| Q2 | 0.76453 | 0.86521 | 0.88597 | 0.88859 | 0.89128 |

**Supplemental Figure 6: Normalization, permutation and cross validation analyses of the metabolomic data. a)** The Liquid chromatography-mass spectroscopy metabolome data before and after normalization. The data was log-transformed and pareto-scaled before any further conducted analyses. **b)** The permutation test statistics at 1000 permutations with observed statistic at $p < 0.001$. The higher number of permutation tests aimed to accurately estimate the statistical significance of the PLS-DA model performance. **c)** Cross-validation (CV) graph showing a 10-fold CV estimate of the predictive ability of our PLS-DA model. R2 shows how well the model predicts the calibration data where Q2 shows how well the model predicts new data. The predictive power of the PLS-DA model dependents on the difference between R2 and Q2 where the smaller the difference, the stronger the predictor is. **d)** Cross-validation calculations of the graph depicted in panel c).

#### Supplemental Videos (MP4 Files)

**Supplemental Videos 1. Multiphoton-second harmonic generation (MP-SHG) bio-imaging examination of the dentin-enamel junction (DEJ) area before and along the course of *ex-vivo* dental caries induction.** The green signals show the MP autofluorescence emitted from the mineralized phase (mainly enamel). The blue signals show the SHG exclusively emitted from the collagen network of dentin. The columns show the maximum intensity projection of the acquired Z-stacks and the 3D renders of sound teeth slices (pre-inoculation), induced incipient caries lesions (caries onset), and induced overt caries lesions. The rows show three representative samples.

**Supplemental Videos 2. Multiphoton-second harmonic generation (MP-SHG) representative Z-stacks of the dentin-enamel junction (DEJ) area of all tested samples after the *ex-vivo* induction of overt caries lesion (17 days).** The green signals show the MP autofluorescence emitted from the mineralized phase (mainly enamel). The blue signals show the SHG exclusively emitted from the collagen network of dentin.

#### Supplemental Tables

**Supplemental Table 1: Composition of the complex Basal Mucin Medium (BMM).**

| Basal Mucin Medium (BMM) | 1L |
|---|---|
| Hog gastric mucin | 2.5 g/L |
| Trypticase peptone | 5.0 g/L |
| Proteose peptone | 10.0 g/L |
| Yeast extract | 5.0 g/L |
| KCl (74.55 g/mol) | 33.5 mmol (2.4975 g) |
| Hemin | 2.5 mg/L |
| Menadione | 0.999mg/L |
| Urea | 60mg/L |
| Arginine | 1.0 mmol |

**Supplemental Table 2: pH measurements of growing biofilms from microcosms without or with sucrose.** For with sucrose microcosms, samples were subjected to sucrose bath pattern 5h/day.

| pH Monitoring | | | |
|---|---|---|---|
| Time After Inoculation | No Sucrose Microcosm (NS) | With Sucrose Microcosm (WS) | |
| | | Sucrose Bath Pattern 5hours/day | |
| | BMM Media | BMM Media 19 hours/day | BMM Media With %5 Sucrose 5 hours/day |
| 0 hour | 6.5-7 | 6.5-7 | 6.5-7 |
| 3 hours | 6.5-7 | 6.5-7 | 6.5-7 |
| 5 hours | 7-8 | 7-8 | 3-4 |
| 10 hours | 7-8 | 7-8 | 3-4 |
| 18 hours | 7-8 | 7-8 | 3-4 |
| 24 hours | 7-8 | 7-8 | 3-4 |
| 30 hours | 7-8 | 7-8 | 3-4 |
| 38 hours | 7-8 | 7-8 | 3-4 |
| 48 hours | 7-8 | 7-8 | 3-4 |
| Media change every 48 hours<br>New inoculum every 96 hours | | | |

**Supplemental Table 3: NanoDrop spectrophotometer results for the DNA yield extracted from the biofilms grown on teeth slices to induce *ex-vivo* dental caries.** (WS, NS) stand for (With Sucrose, No Sucrose) and (T1, T2) stand for (Time Point 1-1st phase/caries onset, Time Point 2-2nd phase/overt lesions).

| No | Sample | Conc. ng/ul | 260/280 | 260/230 |
|---|---|---|---|---|
| ***Time Point 1 (1st phase/caries onset)*** | | | | |
| **1** | 730NS-T1 | 86.8 | 1.89 | 0.69 |
| **2** | 730WS-T1 | 213.3 | 2.1 | 1.39 |
| **3** | 733NS-T1 | 190.2 | 1.9 | 1.6 |
| **4** | 733WS-T1 | 244.38 | 2.0 | 1.95 |
| **5** | 734NS-T1 | 154.8 | 1.93 | 1.44 |
| **6** | 734WS-T1 | 553.02 | 1.98 | 1.65 |
| **7** | 737NS-T1 | 299.94 | 1.89 | 1.54 |
| **8** | 737WS-T1 | 115.2 | 2.0 | 1.19 |
| **9** | 760NS-T1 | 223.5 | 1.98 | 1.73 |
| **10** | 760WS-T1 | 378.78 | 2.0 | 1.84 |
| **11** | 781NS-T1 | 326.58 | 1.89 | 1.48 |
| **12** | 781WS-T1 | 336.18 | 2.15 | 2.16 |
| **13** | 795NS-T1 | 135.12 | 1.79 | 1.18 |
| **14** | 795WS-T1 | 236.5 | 1.99 | 0.92 |
| **15** | 852NS-T1 | 285.6 | 1.7 | 1.24 |
| **16** | 852WS-T1 | 344.64 | 2.0 | 1.98 |
| **17** | 861NS-T1 | 132.78 | 1.89 | 1.48 |
| **18** | 861WS-T1 | 191.16 | 1.99 | 1.78 |
| **19** | 866NS-T1 | 59.1 | 1.97 | 1.80 |
| **20** | 866WS-T1 | 146.16 | 2.3 | 1.49 |
| **21** | 867NS-T1 | 209.16 | 1.97 | 1.56 |
| **22** | 867WS-T1 | 525.8 | 2.0 | 1.82 |
| **23** | ControlToothNS-T1 | 14.8 | 1.78 | 0.53 |
| **24** | ControlToothWS-T1 | 8.6 | 1.84 | 0.51 |
| **25** | MediaNS-T1 | 12.0 | 1.83 | 0.49 |
| **26** | MediaWS-T1 | 9.6 | 1.93 | 0.53 |
| **27** | BALNK-1 | -0.9 | 0.84 | 0.19 |
| **28** | BLANK-2 | -0.3 | 2.25 | 0.03 |
| ***Time Point 2 (2nd phase/overt lesions)*** | | | | |
| **29** | 730NS-T2 | 223.9 | 1.91 | 1.67 |
| **30** | 730WS-T2 | 146.6 | 1.97 | 1.64 |
| **31** | 733NS-T2 | 379.5 | 1.91 | 1.82 |
| **32** | 733WS-T2 | 337.5 | 2.0 | 1.77 |
| **33** | 734NS-T2 | 412.3 | 1.9 | 1.77 |
| **34** | 734WS-T2 | 483.5 | 1.94 | 1.59 |
| **35** | 737NS-T2 | 730.5 | 1.87 | 1.63 |
| **36** | 737WS-T2 | 373.0 | 2.0 | 1.80 |
| **37** | 760NS-T2 | 92.2 | 1.85 | 1.35 |
| **38** | 760WS-T2 | 547.7 | 2.0 | 2.0 |
| **39** | 781NS-T2 | 339.1 | 1.89 | 1.70 |
| **40** | 781WS-T2 | 521.9 | 2.1 | 2.0 |
| **41** | 795NS-T2 | 334.5 | 1.85 | 1.52 |
| **42** | 795WS-T2 | 200.1 | 2.0 | 1.77 |
| **43** | 852NS-T2 | 212.1 | 1.90 | 1.58 |
| **44** | 852WS-T2 | 482.7 | 2.0 | 1.98 |
| **45** | 861NS-T2 | 678.0 | 1.76 | 1.32 |
| **46** | 861WS-T2 | 315.1 | 2.0 | 2.0 |
| **47** | 866NS-T2 | 530.6 | 1.84 | 1.74 |
| **48** | 866WS-T2 | 389.4 | 2.1 | 2.0 |
| **49** | 867NS-T2 | 332.8 | 1.91 | 1.81 |
| **50** | 867WS-T2 | 437.1 | 2.0 | 2.0 |
| **51** | ControlToothNS-T2 | 12.6 | 1.79 | 0.47 |
| **52** | ControlToothWS-T2 | 11.1 | 1.84 | 0.50 |
| **53** | MediaNS-T2 | 12.6 | 1.78 | 0.47 |
| **54** | MediaWS-T2 | 10.6 | 1.79 | 0.49 |
| **55** | BALNK-3 | -0.8 | 0.72 | 0.46 |

**Supplemental Table 4: Mass Spectroscopy (MS) parameters and Selective Reaction Monitoring (SRM) precursor/product ions for the studied metabolites of the central carbon metabolism.**

| Target | Precursor | Product | Declustering Potential (DP) | Collision Energy (CE) |
|---|---|---|---|---|
| Succinate | 117 | 73 | -65 | -16 |
| Fumarate | 115 | 71 | -35 | -14 |
| Malate | 133 | 115 | -50 | -20 |
| Alpha-ketoglutarate | 145 | 101 | -50 | -21 |
| Pyruvate | 87 | 43 | -50 | -12 |
| Lactate | 89 | 43 | -50 | -16 |
| Phosphoenolpyruvate | 166.9 | 79 | -75 | -35 |
| Dihydroxyacetone phosphate | 168.9 | 96.9 | -75 | -50 |
| Glucose 1-phosphate | 259 | 78.8 | -75 | -20 |
| Glucose 6-phosphate | 259 | 199 | -45 | -15 |
| Fructose 6-phosphate | 259 | 97 | -45 | -15 |
| Acetyl coenzyme A | 808 | 461 | -45 | -40 |
| Galactose-1-phosphate | 259 | 241 | -45 | -10 |
| Glyceraldehyde 3-phosphate | 169 | 97 | -45 | -10 |
| Citrate | 191 | 111 | -45 | -10 |
| Fructose 1,6-bisphosphate | 338.9 | 241 | -45 | -10 |
| Ribulose 5-phosphate | 229 | 138.9 | -40 | -20 |
| Ribose 5-phosphate | 229 | 97 | -40 | -20 |

**Supplemental Table 5: Log2 fold change in abundance of taxa associated with dysbiotic (cariogenic) and non-dysbiotic (control) conditions over time (time-based changes).** *q* value denotes the *p* value that has been adjusted for the false discovery rate (FDR) and red asterisks show the statistical significant differences. A negative/positive log2 fold change value indicates the decrease/increase of the corresponding taxa in the first group (Time Point T1-1st phase/caries onset), compared to the second group (Time Point T2-2nd phase/overt lesions), respectively. Key taxa were identified by analysis with DESeq2 differential abundance analysis.

| | No Sucrose (Time Points T1 Vs T2) | log2Fold Change | *p* value | *q* value | With Sucrose (Time Points T1 Vs T2) | log2Fold Change | *p* value | *q* value |
|---|---|---|---|---|---|---|---|---|
| 1 | p__Proteobacteria..c__Gammaproteobacteria..o__Enterobacteriales..f__Enterobacteriaceae.2 | -23.79 | 7.34E-16 | 6.39E-* | p__Firmicutes..c__Bacilli..o__Lactobacillales..f__Enterococcaceae..g__Enterococcus..s__ | -5.13 | 0.0008 | 0.017* |
| 2 | p__Firmicutes..c__Bacilli..o__Lactobacillales..f__Streptococcaceae..g__Streptococcus..s__.1 | 9.43 | 1.82E-05 | 0.0005* | p__Actinobacteria..c__Coriobacteriia..o__Coriobacteriales..f__Coriobacteriaceae..g__Atopobium..s | -14.08 | 1.30E-20 | 5.53E-19* |
| 3 | p__Actinobacteria..c__Coriobacteriia..o__Coriobacteriales..f__Coriobacteriaceae..g__Atopobium..s__ | -10.098 | 1.10E-06 | 4.77E-05* | p__Actinobacteria..c__Actinobacteria..o__Actinomycetales..f__Corynebacteriaceae..g__Corynebacterium | -27.05 | 3.78E-25 | 3.21E-23* |
| 4 | p__Firmicutes..c__Bacilli..o__Lactobacillales..f__Enterococcaceae..g__Enterococcus..s__ | -3.452 | 0.0563 | 0.544 | p__Firmicutes..c__Bacilli..o__Bacillales..f__Bacillaceae..g__Bacillus.1 | 21.66 | 2.12E-13 | 6.01E-12* |
| 5 | p__Proteobacteria..c__Gammaproteobacteria..o__Pasteurellales..f__Pasteurellaceae..g__Haemophilus..s__parainfluenzae | 3.57 | 0.0258 | 0.331 | p__Proteobacteria..c__Gammaproteobacteria..o__Enterobacteriales..f__Enterobacteriaceae | -4.475 | 0.0058 | 0.082 |
| 6 | p__Firmicutes..c__Bacilli..o__Lactobacillales..f__Streptococcaceae..g__Streptococcus..s__ | 5.68 | 0.035 | 0.390 | p__Firmicutes..c__Bacilli..o__Bacillales..f__Bacillaceae..g__Bacillus | 1.634 | 0.033 | 0.180 |
| 7 | p__Fusobacteria..c__Fusobacteriia..o__Fusobacteriales..f__Fusobacteriaceae..g__Fusobacterium..s__.1 | 3.91 | 0.026 | 0.331 | p__Firmicutes..c__Clostridia..o__Clostridiales..f__Veillonellaceae..g__Veillonella..s__dispar | -2.58 | 0.033 | 0.180 |
| 8 | p__Firmicutes..c__Clostridia..o__Clostridiales..f__Veillonellaceae..g__Veillonella..s__parvula.1 | -7.98 | 0.006 | 0.135 | p__Firmicutes..c__Bacilli..o__Lactobacillales..f__Lactobacillaceae..g__Lactobacillus..s__.1 | -4.60 | 0.0172 | 0.1330 |
| 9 | p__Bacteroidetes..c__Bacteroidia..o__Bacteroidales..f__.Paraprevotellaceae...g__.Prevotella...s__.6 | 6.625 | 0.024 | 0.331 | p__Proteobacteria..c__Gammaproteobacteria..o__Pasteurellales..f__Pasteurellaceae..g__Haemophilus..s__parainfluenzae | 4.29 | 0.009 | 0.1023 |
| 10 | | | | | p__Fusobacteria..c__Fusobacteriia..o__Fusobacteriales..f__Fusobacteriaceae..g__Fusobacterium..s__ | -3.56 | 0.0402 | 0.180 |
| 11 | | | | | p__Firmicutes..c__Bacilli..o__Lactobacillales..f__Carnobacteriaceae..g__Granulicatella..s__ | -1.50 | 0.037 | 0.18007 |
| 12 | | | | | p__Firmicutes..c__Clostridia..o__Clostridiales..f__.Tissierellaceae...g__Parvimonas..s__ | -4.99 | 0.006 | 0.08304 |
| 13 | | | | | p__Firmicutes..c__Bacilli..o__Gemellales..f__Gemellaceae | -2.96 | 0.0126 | 0.10722 |
| 14 | | | | | p__Firmicutes..c__Clostridia..o__Clostridiales..f__Veillonellaceae..g__Veillonella..s__dispar.3 | -7.36 | 0.0109 | 0.10314 |
| 15 | | | | | p__Firmicutes..c__Bacilli..o__Bacillales..f__Planococcaceae..g__Lysinibacillus | -6.16 | 0.0375 | 0.1800 |
| 16 | | | | | p__Proteobacteria..c__Betaproteobacteria..o__Neisseriales..f__Neisseriaceae..g__Eikenella..s__.1 | -8.39 | 0.004 | 0.07621 |
| 17 | | | | | p__Actinobacteria..c__Actinobacteria..o__Actinomycetales..f__Actinomycetaceae..g__Actinomyces..s__.2 | -6.20 | 0.036 | 0.1800 |
| 18 | | | | | p__Firmicutes..c__Clostridia..o__Clostridiales..f__Veillonellaceae..g__Megasphaera..s__.2 | -6.87 | 0.0202 | 0.1431 |
| 19 | | | | | p__Firmicutes..c__Clostridia..o__Clostridiales..f__Lachnospiraceae..g__Moryella..s_ | -6.14 | 0.0381 | 0.1800 |

**Supplemental Table 6: Log2 fold change in abundance of taxa associated with each time point per treatment (treatment-based changes).** *q* value denotes the *p* value that has been adjusted for the false discovery rate (FDR) and red asterisks show the statistical significant differences. A negative/positive log2 fold change value indicates the decrease/increase of the corresponding taxa in the first group (No Sucrose "NS") compared to the second group ( With Sucrose "WS"), respectively. (TimePoint1 stands for the 1$^{st}$ phase/caries onset and TimePoint2 stands for 2$^{nd}$ phase/overt lesions). Key taxa were identified by analysis with DESeq2 differential abundance analysis.

| | TimePoint1 (Treatment NS Vs WS) | log2Fold Change | *p* value | *q* value | TimePoint2 (Treatment NS Vs WS) | log2Fold Change | *p* value | *q* value |
|---|---|---|---|---|---|---|---|---|
| 1 | p__Firmicutes..c__Bacilli..o__Lactobacillales..f__Lactobacillaceae..g__Lactobacillus..s__.1 | -25.07 | 1.89E-17 | 2.36E-16* | p__Firmicutes..c__Bacilli..o__Bacillales..f__Bacillaceae..g__Bacillus | 2.25 | 0.005 | 0.027* |
| 2 | p__Proteobacteria..c__Gammaproteobacteria..o__Enterobacteriales..f__Enterobacteriaceae.2 | -27.2 | 2.28E-20 | 3.80E-19* | p__Firmicutes..c__Bacilli..o__Lactobacillales..f__Enterococcaceae..g__Enterococcus..s__ | -3.67 | 1.37E-06 | 1.05E-05* |
| 3 | p__Firmicutes..c__Clostridia..o__Clostridiales..f__Veillonellaceae..g__Veillonella..s__dispar.1 | -24.54 | 1.03E-22 | 5.15E-21* | p__Firmicutes..c__Bacilli..o__Lactobacillales..f__Lactobacillaceae..g__Lactobacillus..s__.1 | -12.90 | 7.47E-22 | 2.02E-20* |
| 4 | p__Firmicutes..c__Clostridia..o__Clostridiales..f__Veillonellaceae..g__Veillonella..s__parvula.1 | -21.74 | 1.76E-13 | 1.76E-12* | p__Firmicutes..c__Bacilli..o__Lactobacillales..f__Streptococcaceae..g__Streptococcus..s__ | -13.53 | 1.49E-16 | 1.61E-15* |
| 5 | p__Bacteroidetes..c__Bacteroidia..o__Bacteroidales..f__Porphyromonadaceae..g__Porphyromonas..s__ | 8.509 | 4.94E-05 | 0.00026* | p__Firmicutes..c__Bacilli..o__Lactobacillales..f__Streptococcaceae..g__Streptococcus..s__.1 | -11.67 | 1.05E-10 | 9.41E-10* |
| 6 | p__Firmicutes..c__Bacilli..o__Lactobacillales..f__Streptococcaceae..g__Streptococcus..s__.5 | 8.72 | 0.0003439 | 0.0017* | p__Actinobacteria..c__Coriobacteriia..o__Coriobacteriales..f__Coriobacteriaceae..g__Atopobium..s__ | -3.84 | 0.002 | 0.012* |
| 7 | p__Firmicutes..c__Bacilli..o__Lactobacillales..f__Aerococcaceae..g__Abiotrophia..s__ | 9.4190 | 1.19E-10 | 9.91E-10* | p__Firmicutes..c__Clostridia..o__Clostridiales..f__Veillonellaceae..g__Veillonella..s__dispar.1 | -27.35 | 3.12E-30 | 1.69E-28* |
| 8 | p__Actinobacteria..c__Actinobacteria..o__Actinomycetales..f__Actinomycetaceae..g__Actinomyces..s__.2 | 8.373 | 3.75E-05 | 0.00023* | p__Firmicutes..c__Bacilli..o__Bacillales..f__Staphylococcaceae..g__Staphylococcus | -7.46 | 0.0020 | 0.012* |
| 9 | p__TM7..c__TM7.3..o__CW040..f__..g__..s__ | 24.767 | 1.18E-20 | 2.94E-19* | p__Bacteroidetes..c__Bacteroidia..o__Bacteroidales..f__Porphyromonadaceae..g__Porphyromonas..s | 4.53 | 0.002 | 0.0479* |
| 10 | p__Firmicutes..c__Bacilli..o__Bacillales..f__Paenibacillaceae..g__Paenibacillus..s__.1 | 7.596 | 2.22E-05 | 0.0001* | p__Firmicutes..c__Bacilli..o__Bacillales..f__Paenibacillaceae..g__Paenibacillus..s__ | 3.0161 | 0.0113 | 0.047* |
| 11 | p__Firmicutes..c__Bacilli..o__Bacillales..f__Paenibacillaceae..g__Paenibacillus..s__.2 | 7.391 | 0.00052 | 0.0023* | p__Firmicutes..c__Bacilli..o__Lactobacillales..f__Streptococcaceae..g__Streptococcus..s__anginosus | -26.26 | 2.21E-21 | 3.97E-20* |
| 12 | p__Firmicutes..c__Bacilli..o__Lactobacillales..f__Streptococcaceae..g__Streptococcus..s__ | -4.96192 | 0.0024 | 0.010015* | p__Fusobacteria..c__Fusobacteriia..o__Fusobacteriales..f__Fusobacteriaceae..g__Fusobacterium..s__.7 | -25.24 | 1.16E-17 | 1.57E-16* |
| 13 | p__Firmicutes..c__Bacilli..o__Lactobacillales..f__Carnobacteriaceae..g__Granulicatella..s__ | 2.057081 | 0.0146 | 0.048677* | p__Proteobacteria..c__Betaproteobacteria..o__Neisseriales..f__Neisseriaceae..g__Eikenella..s__.1 | -7.602 | 0.0100 | 0.047* |
| 14 | p__Firmicutes..c__Bacilli..o__Lactobacillales..f__Streptococcaceae..g__Streptococcus..s__.1 | -3.408490 | 0.0100 | 0.038577* | k__Bacteria..p__Bacteroidetes..c__Bacteroidia..o__Bacteroidales..f__Porphyromonadaceae..g__Porphyromonas..s__ | 4.539 | 0.0115 | 0.0475* |
| 15 | p__Firmicutes..c__Bacilli..o__Gemellales..f__Gemellaceae | 3.355926 | 0.011 | 0.042177* | p__Firmicutes..c__Bacilli..o__Lactobacillales..f__Streptococcaceae..g__Streptococcus | -2.4409 | 0.0296 | 0.09407 |
| 16 | p__Firmicutes..c__Bacilli..o__Bacillales..f__Staphylococcaceae..g__Staphylococcus | -5.48943 | 0.01809 | 0.055311* | p__Firmicutes..c__Bacilli..o__Lactobacillales..f__Streptococcaceae..g__Streptococcus..s__.5 | 6.12373 | 0.0218 | 0.07848 |
| 17 | p__Firmicutes..c__Bacilli..o__Bacillales..f__Paenibacillaceae..g__Paenibacillus..s__ | 3.200182 | 0.0188 | 0.055311* | p__Firmicutes..c__Bacilli..o__Bacillales..f__Paenibacillaceae..g__Paenibacillus..s__.1 | 4.6595 | 0.0200 | 0.07749 |
| 18 | p__Proteobacteria..c__Epsilonproteobacteria..o__Campylobacterales..f__Campylobacteraceae..g__Campylobacter..s__.1 | 5.879356 | 0.04 | 0.100744 | p__Firmicutes..c__Bacilli..o__Bacillales..f__Paenibacillaceae..g__Paenibacillus..s__.2 | 4.83977 | 0.0275 | 0.09285 |
| 19 | p__Fusobacteria..c__Fusobacteriia..o__Fusobacteriales..f__Fusobacteriaceae..g__Fusobacterium..s__.7 | -5.908617 | 0.04 | 0.103253 | | | | |
| 20 | p__Bacteroidetes..c__Bacteroidia..o__Bacteroidales..f__.Paraprevotellaceae...g__.Prevotella...s__.6__ | 6.46963 | 0.028 | 0.074817 | | | | |
| 21 | p__Proteobacteria..c__Epsilonproteobacteria..o__Campylobacterales..f__Campylobacteraceae..g__Campylobacter..s__.2 | 5.78531 | 0.05 | 0.109037 | | | | |
| 22 | p__Fusobacteria..c__Fusobacteriia..o__Fusobacteriales..f__Fusobacteriaceae..g__Fusobacterium..s__.10 | 5.191306 | 0.07 | 0.15802 | | | | |
| 23 | p__Bacteroidetes..c__Flavobacteriia..o__Flavobacteriales..f__Flavobacteriaceae..g__Capnocytophaga..s__ochracea | 6.59149 | 0.025 | 0.07102 | | | | |
| 24 | p__Firmicutes..c__Bacilli..o__Lactobacillales..f__Streptococcaceae..g__Streptococcus..s__.8 | 6.24084 | 0.0345 | 0.08638 | | | | |
| 25 | p__Firmicutes..c__Bacilli..o__Lactobacillales..f__Streptococcaceae..g__Streptococcus__ | -1.571016 | 0.0561 | 0.117043 | | | | |

**Supplemental Table 7: Indicator Value "IndVal" index for species analysis showing taxa associated with particular groups.** "NS_T1" stands for No Sucrose at Time Point 1-1st phase/caries onset, "NS_T2" stands for No Sucrose at Time Point 2-2nd phase/overt lesions, "WS_T1" stands for With Sucrose at Time Point 1-1st phase/caries onset and "WS_T2" stands for With Sucrose at Time Point 2-2nd phase/over lesions.

| Amplicon Sequence Variant (ASV) | Cluster | Indicator | Probability |
|---|---|---|---|
| p__Firmicutes..c__Bacilli..o__Lactobacillales..f__Aerococcaceae..g__Abiotrophia..s__ | NS_T1 | 0.8056 | 0.001 |
| p__Bacteroidetes..c__Bacteroidia..o__Bacteroidales..f__Porphyromonadaceae..g__Porphyromonas..s__ | NS_T1 | 0.6047 | 0.002 |
| p__Proteobacteria..c__Gammaproteobacteria..o__Pasteurellales..f__Pasteurellaceae..g__Haemophilus..s__parainfluenzae | NS_T1 | 0.5823 | 0.003 |
| p__Fusobacteria..c__Fusobacteriia..o__Fusobacteriales..f__Fusobacteriaceae..g__Fusobacterium..s__.1 | NS_T1 | 0.5739 | 0.004 |
| p__Firmicutes..c__Bacilli..o__Bacillales..f__Paenibacillaceae..g__Paenibacillus..s__.1 | NS_T1 | 0.5635 | 0.002 |
| p__Firmicutes..c__Bacilli..o__Lactobacillales..f__Carnobacteriaceae..g__Granulicatella..s__ | NS_T1 | 0.5116 | 0.001 |
| p__Firmicutes..c__Bacilli..o__Bacillales..f__Paenibacillaceae..g__Paenibacillus..s__ | NS_T1 | 0.5109 | 0.011 |
| p__Actinobacteria..c__Actinobacteria..o__Actinomycetales..f__Actinomycetaceae..g__Actinomyces..s__.2 | NS_T1 | 0.4933 | 0.013 |
| p__Firmicutes..c__Bacilli..o__Lactobacillales..f__Streptococcaceae..g__Streptococcus..s__.3 | NS_T1 | 0.4755 | 0.028 |
| p__Firmicutes..c__Bacilli..o__Gemellales..f__Gemellaceae | NS_T1 | 0.4713 | 0.013 |
| p__Firmicutes..c__Bacilli..o__Bacillales..f__Paenibacillaceae..g__Paenibacillus..s__.2 | NS_T1 | 0.4101 | 0.013 |
| p__TM7..c__TM7.3..o__CW040..f__..g__..s__ | NS_T1 | 0.4009 | 0.018 |
| p__Firmicutes..c__Bacilli..o__Lactobacillales..f__Streptococcaceae..g__Streptococcus..s__.5 | NS_T1 | 0.387 | 0.01 |
| p__Firmicutes..c__Bacilli..o__Bacillales..f__Bacillaceae..g__Bacillus.1 | NS_T1 | 0.3819 | 0.016 |
| p__Bacteroidetes..c__Flavobacteriia..o__Flavobacteriales..f__.Weeksellaceae...g__..s__.1 | NS_T1 | 0.2987 | 0.044 |
| p__Firmicutes..c__Clostridia..o__Clostridiales..f__.Tissierellaceae...g__Parvimonas..s__ | NS_T2 | 0.5704 | 0.001 |
| p__Proteobacteria..c__Gammaproteobacteria..o__Enterobacteriales..f__Enterobacteriaceae | NS_T2 | 0.465 | 0.002 |
| p__Proteobacteria..c__Betaproteobacteria..o__Neisseriales..f__Neisseriaceae..g__Eikenella..s__ | NS_T2 | 0.4564 | 0.015 |
| p__Firmicutes..c__Bacilli..o__Lactobacillales..f__Streptococcaceae..g__Streptococcus..s__.1 | WS_T1 | 0.6698 | 0.001 |
| p__Firmicutes..c__Bacilli..o__Lactobacillales..f__Streptococcaceae..g__Streptococcus | WS_T1 | 0.6665 | 0.001 |
| p__Firmicutes..c__Bacilli..o__Lactobacillales..f__Streptococcaceae..g__Streptococcus..s__ | WS_T1 | 0.6507 | 0.002 |
| p__Firmicutes..c__Bacilli..o__Lactobacillales..f__Lactobacillaceae..g__Lactobacillus..s__.1 | WS_T2 | 0.8546 | 0.001 |
| p__Actinobacteria..c__Coriobacteriia..o__Coriobacteriales..f__Coriobacteriaceae..g__Atopobium..s__ | WS_T2 | 0.8077 | 0.001 |
| p__Firmicutes..c__Bacilli..o__Lactobacillales..f__Enterococcaceae..g__Enterococcus..s__ | WS_T2 | 0.6441 | 0.001 |
| p__Firmicutes..c__Bacilli..o__Lactobacillales..f__Streptococcaceae..g__Streptococcus..s__anginosus | WS_T2 | 0.3194 | 0.029 |
